# An integrated model system to gain mechanistic insights into biofilm formation and antimicrobial resistance development in *Pseudomonas aeruginosa* MPAO1

**DOI:** 10.1101/2020.02.06.936690

**Authors:** Adithi R. Varadarajan, Raymond N. Allan, Jules D. P. Valentin, Olga E. Castañeda Ocampo, Vincent Somerville, Franziska Pietsch, Matthias T. Buhmann, Jonathan West, Paul J. Skipp, Henny C. van der Mei, Qun Ren, Frank Schreiber, Jeremy S. Webb, Christian H. Ahrens

## Abstract

*Pseudomonas aeruginosa* MPAO1 is the parental strain of the widely utilized transposon mutant collection for this important clinical pathogen. Here, we validate a model system to identify genes involved in biofilm growth and antibiotic resistance.

Our model employs a genomics-driven workflow to assemble the complete MPAO1 genome, identify unique and conserved genes by comparative genomics with the PAO1 reference strain and missed genes by proteogenomics. Among over 200 unique MPAO1 genes, we identified six general essential genes that were overlooked when mapping public Tn-seq datasets against PAO1, including an antitoxin. Genomic data were integrated with phenotypic data from an experimental workflow using a user-friendly, soft lithography-based microfluidic flow chamber for biofilm growth. Experiments conducted across three laboratories delivered reproducible data on *P. aeruginosa* biofilms and validated both known and novel genes involved in biofilm growth and antibiotic resistance identified in screens of the mutant collection. Differential protein expression data from planktonic cells versus biofilm confirmed upregulation of candidates known to affect biofilm formation, of structural and secreted proteins of type six secretion systems, and provided proteogenomic evidence for some missed MPAO1 genes. This integrated, broadly applicable model promises to improve the mechanistic understanding of biofilm formation, antimicrobial tolerance and resistance evolution.

## Introduction

*Pseudomonas aeruginosa* is a gram-negative bacterium ubiquitously present in soil, water and different animal hosts [1]. As an opportunistic human pathogen [2] it can cause sepsis, and chronic wound and lung infections (i.e., cystic fibrosis), especially in immunocompromised individuals. Two mechanisms complicate the treatment of *P. aeruginosa* infections. It forms recalcitrant biofilms in which the bacterial cells have an increased tolerance against antimicrobial compounds [3, 4]. In addition, worldwide, multiple genetic variants have acquired antimicrobial resistance (AMR) traits [5], either through acquisition of resistance genes on mobile genetic elements such as plasmids [6] or through *de novo* mutations of chromosomal genes [7]. Furthermore, mutations affecting outer membrane porins and multi-drug efflux pumps can mediate resistance to almost all major antibiotic classes and several important biocides [8, 9]. *P. aeruginosa* thus also belongs to the notorious group of ESKAPE pathogens, which represent the lead cause of worldwide nosocomial infections (*Enterococcus faecium, Staphylococcus aureus, Klebsiella pneumoniae, Acinetobacter baumannii, P. aeruginosa*, and *Enterobacter* species) [10, 11]. Clinically most relevant are the resistances of *P. aeruginosa* strains against fluoroquinolones, aminoglycosides and beta-lactams, and against the last-resort antibiotic colistin (a polymyxin). In 2017, the World Health Organization (WHO) classified carbapenem-resistant *P. aeruginosa* strains in the highest priority group of “critical pathogens”. New treatment options informed by a more detailed molecular understanding of how and why resistance emerges during treatment, and how resistance is transmitted, are urgently needed for such critical pathogens.

Increased antimicrobial tolerance, a fundamental property of biofilms [12] is well-studied [13] and four mechanisms play a major role: (i) under nutrient-limited conditions in biofilms, *P. aeruginosa* expresses phenotypic variants, i.e., dormant cells that are less susceptible to antibiotics which target actively dividing cells [14]; (ii) *P. aeruginosa* form a protective extra-cellular matrix composed of polysaccharides, proteins and DNA that limits the diffusion of antimicrobial substances or sequesters them, such that biofilm cells experience a decreased antimicrobial dosage [15]; (iii) anoxic conditions exist within the biofilm limiting the efficacy of antibiotics that require aerobic metabolic activity and the generation of reactive oxygen species [16]; (iv) sub-inhibitory concentrations of antibiotics induce increased rates of mutation, recombination and lateral transfer. The mutation rate in biofilms has been reported to be up to 100 times higher than in planktonic cells [17], significantly accelerating the development of antibiotic resistant mutants. Together, these mechanisms lead to hard-to-treat, chronic infections during which *P. aeruginosa* can persist and further evolve within the host in the presence of antimicrobial substances. Evolution within biofilms is highly parallel and differs significantly from evolution of planktonic cells [18]. However, the evolutionary drivers of within-biofilm AMR evolution remain poorly understood. Their study requires well-defined model systems and tools, including model strains with complete genomic background information, genetic tools and flow chambers allowing representative and reproducible growth of *P. aeruginosa* biofilms.

The canonical reference model strain for *P. aeruginosa* is PAO1, also referred to as PAO1-UW. Its complete genome sequence was published in 2000 [2], which allowed many breakthrough discoveries. However, a number of closely related PAO1 strains exist that differ in their phenotypic appearances [19]. These include *P. aeruginosa* strain MPAO1 [20], the parental strain of the widely utilized transposon insertion mutant library from the University of Washington (UW) [21]. Such mutant collections represent highly valuable resources to uncover new functions and essential genes in genome-wide screens [21], for example genes relevant for resistance against certain antibiotics [22] [23]. They have also been used to define general essential genes, which are essential under more than one relevant growth condition [24, 25], and a subset of the core essential genes *of P. aeruginosa* PAO1 and PA14 were shown to exhibit differential essentiality [26]. However, the utility of such libraries to identify gain of function mutations is limited and polar effects need to be controlled for [27]. Notably, no complete MPAO1 genome sequence was available. Improvements in next generation sequencing (NGS) technologies [28] and assembly algorithms nowadays allow researchers to readily generate complete *de novo* genome assemblies for most prokaryotes except a few percent of strains with highly complex repeat regions [29]. Such fully resolved genomes are advantageous compared to fragmented short read-based genome assemblies that can sometimes even miss conserved core genes [30]; they are an ideal basis for subsequent functional genomics and systems biology studies, allowing to identify novel genes missed in genome annotations by proteogenomics [31]. As high-quality reference points, complete genomes also enable the identification of genes important for the evolutionary adaptation of *P. aeruginosa* in biofilms exposed to antimicrobial substances by deep sequencing [18].

Here, we set out to develop, validate and make available to the community a model system to study the biofilm-associated adaptation to antimicrobials and AMR evolution in *P. aeruginosa* MPAO1. Conceptually, the model was designed to integrate genotype information with phenotypic data and to leverage the valuable genetic tools and wealth of functional genomics datasets that exist for important bacterial model organisms. Important elements include the complete MPAO1 genome sequence and the design for a standardized flow chamber based on accessible soft lithography replication in poly(dimethylsiloxane) (PDMS) that can deliver laminar flow conditions relevant to typical biofilm niches. Comparative genomics with the PAO1-UW reference strain uncovered numerous MPAO1-unique genes. Strikingly, these included 39 essential genes that had been missed so far by performing reference-based mapping of public Tn-seq datasets. Proof of principle experiments highlighted reproducible biofilm growth using the microfluidic flow chamber, and identified both known and novel genes important for biofilm growth and AMR through microtiter plate screening of the mutant library. A differential (planktonic vs. biofilm) proteomic dataset uncovered genes known to play a role in biofilm formation. Finally, a publicly available, integrated proteogenomics search database enables identification of novel genes in MPAO1.

## Results

### *De novo* genome assembly of MPAO1

The availability of a complete genome sequence is an important pre-requisite to study the evolution of resistance to antimicrobials in biofilms. An analysis of more than 9,300 completely sequenced and publicly available bacterial genomes [29] (see Methods) indicated that they comprised 106 *P. aeruginosa* strains. However, only two *P. aeruginosa* PAO1 strains were among these, including the PAO1 type strain (Genbank AE004091), also called PAO1-UW [2]. In contrast, the only strain annotated as MPAO1, i.e. the founder strain of the transposon mutant library available from the UW [21], had been sequenced with Illumina short reads, assembled into 140 contigs [32] and deposited in the Pseudomonas genome DB (http://www.pseudomonas.com/strain/show?id=659; Genbank ASM24743v2) [33]. To provide an optimal basis for subsequent functional genomics and evolution studies for *P. aeruginosa* strain MPAO1, we thus first sequenced and *de novo* assembled its complete genome. Due to the genomic differences reported for MPAO1 and PAO1 [20] and the fact that many of the 106 completely sequenced *P. aeruginosa* strains have difficult to assemble genomes with long repeat pairs in excess of 10 kilobases (kb) (38/106), so-called class III genomes [29], we used third generation long reads from Pacific Biosciences’ (PacBio) RSII platform. By relying on size-selected fragments (average length 9 kb; see Methods), a single bacterial chromosome could be assembled. Additional genome polishing steps with Illumina MiSeq data (300bp, PE reads) allowed to remove remaining homopolymer errors in the PacBio assembly [34]. The final, high-quality MPAO1 genome consisted of one chromosome of 6,275,467 bp and coded for 5,926 genes (Genbank CP027857; **Table 1**). An overview of selected predicted genome features (see Methods) is shown in Supplementary **Table 1**. To facilitate data mining and comparison, we also provide an extensive annotation of all 5,799 protein-coding genes. This includes information on conserved and MPAO1-unique genes compared to PAO1, the respective reciprocal best BLAST hits, protein domains, families, Gene Ontology (GO) classification, predictions of subcellular localization, lipoproteins, secreted and described membrane-localized proteins, as well as gene essentiality status and protein expression data below (Supplementary **Table 2**).

**Table 1.**
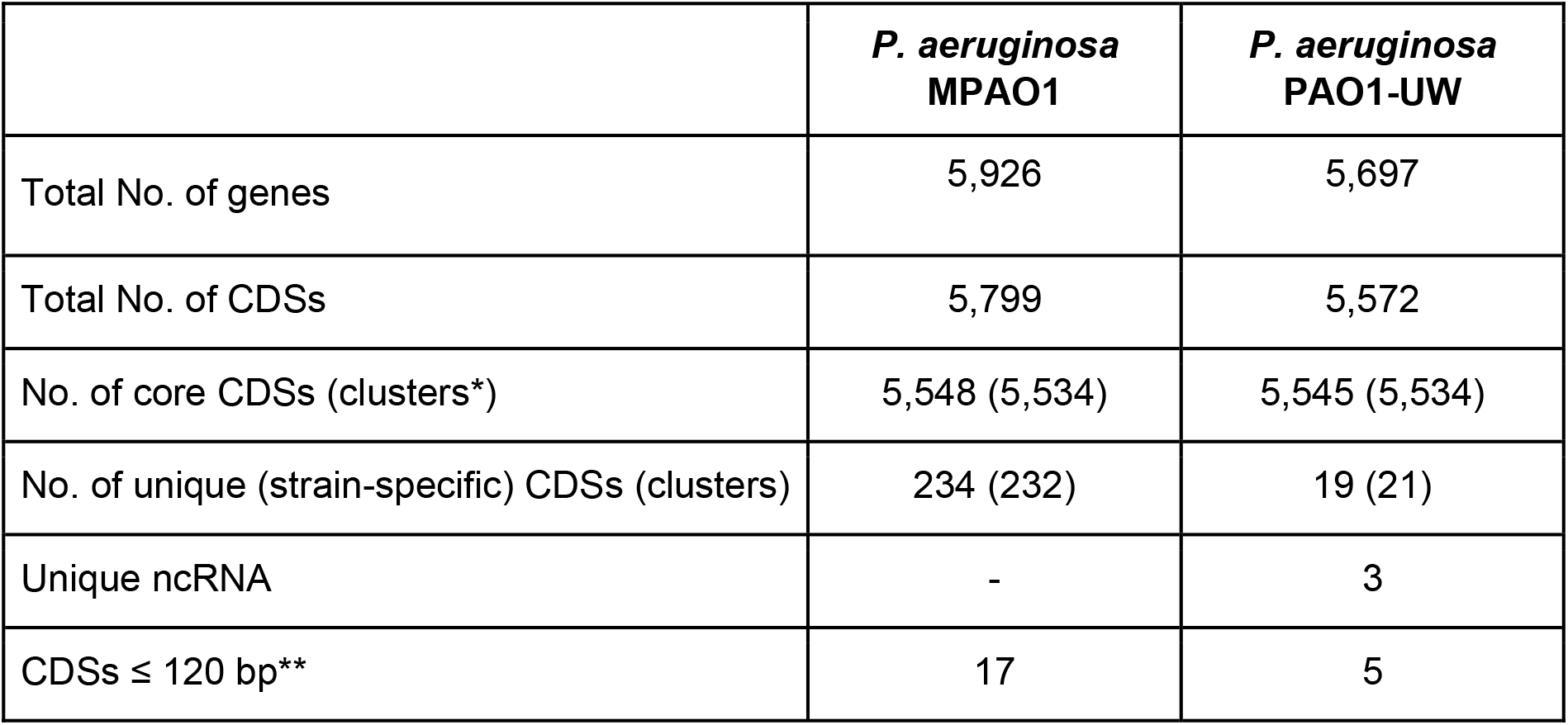
Summary over core and strain-specific CDS of strains MPAO1 and PAO1-UW. *All individual CDS are shown including those that are grouped in gene clusters (paralogs) in Fig. 1c. **CDS of 120 bp or below are not considered (see Methods).

**Table 2.** List of 18 selected MPAO1-unique genes along with their essentiality classification in all 16 Tn-seq samples [24] and comments about their genomic location. Information about all MPAO1-unique essential genes is available in Supplementary **Table 7**.

| Locus | Gene annotation | General essential | Essential in x/16 samples | Comment |
| --- | --- | --- | --- | --- |
| MPAO1_22380 | type II toxin-antitoxin system Phd/YefM family antitoxin | yes | 16 | Prophage 2 |
| MPAO1_00215 | hypothetical protein | yes | 15 | *Operon? |
| MPAO1_10410 | hypothetical protein | yes | 14 |  |
| MPAO1_22450 | DNA-binding protein | yes | 14 | Prophage 2 |
| MPAO1_25260 | cytidine deaminase |  | 12 |  |
| MPAO1_12950 | hypothetical protein | yes | 11 |  |
| MPAO1_00230 | hypothetical protein |  | 10 | *Operon? |
| MPAO1_20095 | hypothetical protein |  | 10 |  |
| MPAO1_02335 | dihydropyrimidinase |  | 9 |  |
| MPAO1_15010 | 6-O-methylguanine DNA methyltransferase |  | 9 |  |
| MPAO1_15215 | amino acid permease |  | 9 |  |
| MPAO1_18025 | ferredoxin |  | 9 |  |
| MPAO1_02315 | oxidoreductase |  | 8 |  |
| MPAO1_05695 | hypothetical protein | yes | 8 | Bacteriocin (GO) |
| MPAO1_08710 | DUF3304 domain-containing protein |  | 8 |  |
| MPAO1_10195 | universal stress protein |  | 8 |  |
| MPAO1_14380 | glycosyltransferase |  | 8 |  |
| MPAO1_24865 | hypothetical protein |  | 8 | Prophage 3 |

### Comparative genomics of MPAO1 and PAO1 strains

An alignment of our *de novo* assembled MPAO1 genome with that of the MPAO1/P1 strain [32] revealed that overall, 42,813 bp of our complete genome sequence were missed by the 140 contigs of the available fragmented Illumina assembly (**Fig. 1a**). This comprised 66 genes (52 protein coding genes, (CDS)) either missed completely or partially, including eight of 12 rRNA genes (75%) and six of 63 tRNA genes (11%). Among the CDS, the essential gene *ftsY* encoding the signal recognition particle-docking protein FtsY was missing, four of eight (50%) non-ribosomal peptide synthetase (NRPS) genes, three of six (50%) filamentous hemagglutinin N-terminal domain protein coding genes and three of 10 (30%) type VI secretion system (T6SS) VgrG effector proteins (Supplementary **Table 2**). The analysis of the number of interrupted genes or pseudogenes also confirmed the fragmented nature of the MPAO1/P1 genome compared to the complete genomes of both our assembly and the PAO1-UW type strain (Supplementary **Fig. 1**). Importantly, a key study of the genotypic and phenotypic diversity of *P. aeruginosa* PAO1 strains recently reported 10 PAO1/MPAO1 laboratory isolates as complete genomes [19]. As all 10 genomes have been assembled using Illumina data into sets of contigs, strictly speaking, they are not fully assembled, closed genome sequences. Indeed, the genomes of the two MPAO1 strains in that list (PAO1-2017-E, 71 contigs, whole genome shotgun (WGS) QZGA00000000 and PAO1-2017-I, 70 contigs, WGS QZGE00000000) also lacked a similar amount of genomic sequence (56.5 and 59.4 kb) and number of genes (55, 62) or CDS (40, 47) respectively, compared to our complete genome (Supplementary **Table 2**).

**Fig. 1.**
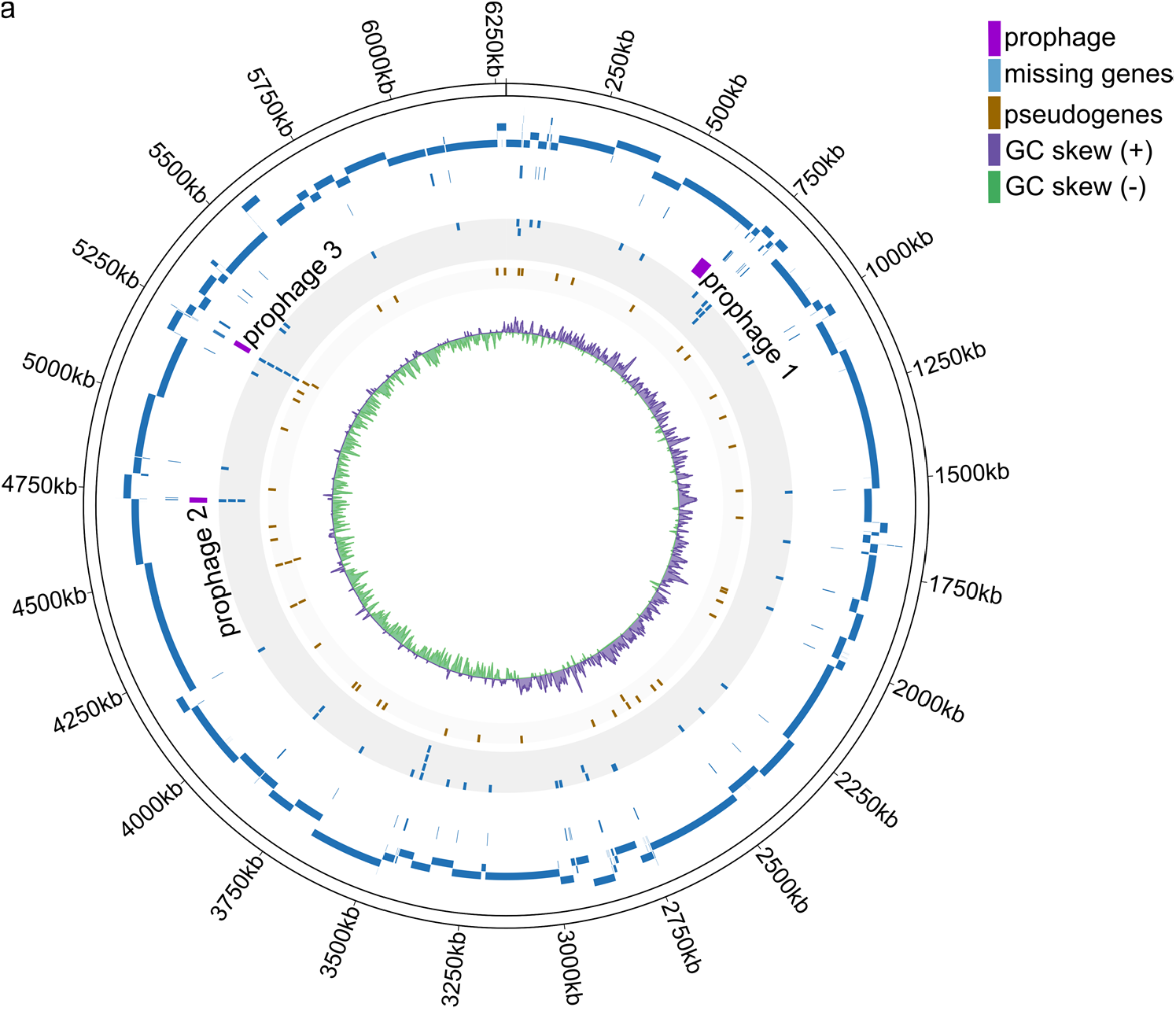
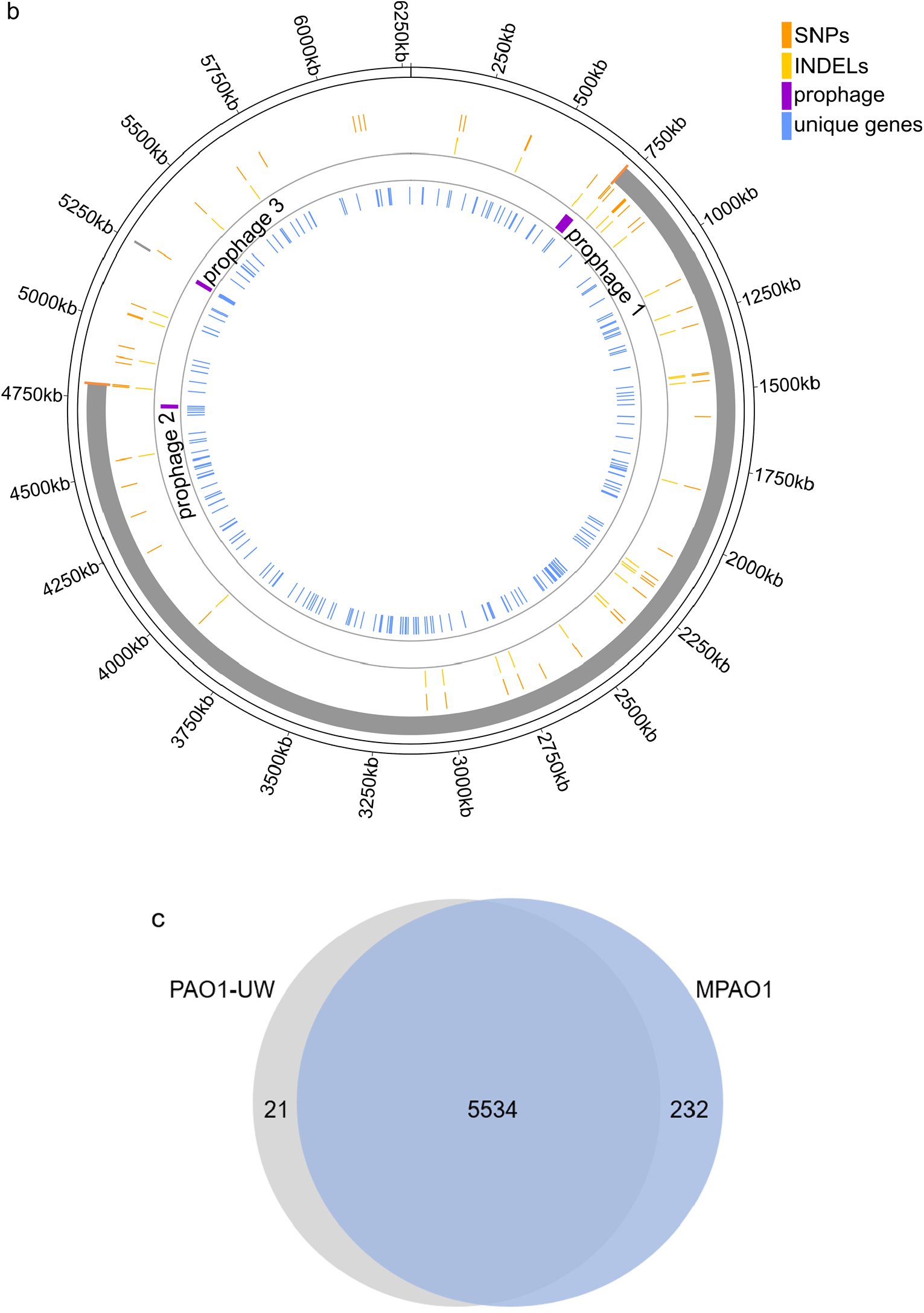
Genome map of *P. aeruginosa* MPAO1 and comparison to other strains. (**a**) The Circos plot visualizes the comparison of our complete MPAO1 genome (outer circle with genome coordinates) and that of strain MPAO1/P1 (second circle; blue), the respective gaps (third circle; blue) followed by annotated prophages (fourth circle; purple), missing genes (fifth circle, light blue), pseudogenes (sixth circle; brown), and GC skew (seventh circle; positive - purple; negative - green). (**b**) Differences of the MPAO1 genome compared to the PAO1 reference strain. Going from outer towards inner circles, the following genome features are shown: (1) a large inversion (gray) flanked by rRNAs (not shown), (2) SNPs (dark orange), (3) INDELs (light orange) (4) prophages (purple), (5) genes unique to MPAO1 (blue). (**c**) Comparative genomic analysis of *P. aeruginosa* strains MPAO1 and PAO1-UW. The Venn diagram shows the core gene clusters (paralogous genes are grouped into the same cluster provided they belong to a syntenic genomic region) and the respective number of strain-specific CDS clusters.

Next, to explore the extent of strain-specific genomic differences, we created an alignment of our *de novo* assembled MPAO1 genome with that of *P. aeruginosa* strain PAO1-UW. This analysis confirmed the major differences reported previously [20], i.e. the presence of a third prophage region (12.8 kb, 20 genes; genome coordinates 5,241,813 - 5,254,613) in strain MPAO1 (**Fig. 1b**) and the absence of a ∼1 kb genome fragment (leading to a pseudogene annotation for MPAO1_24940 in MPAO1). An analysis of smaller differences between the genomes confirmed the 16 SNPs reported previously [20], and identified 176 additional SNPs and INDELs between MPAO1 and PAO1 that had not been reported by Klockgether and colleagues [20] (Supplementary **Table 3**).

**Table 3.**
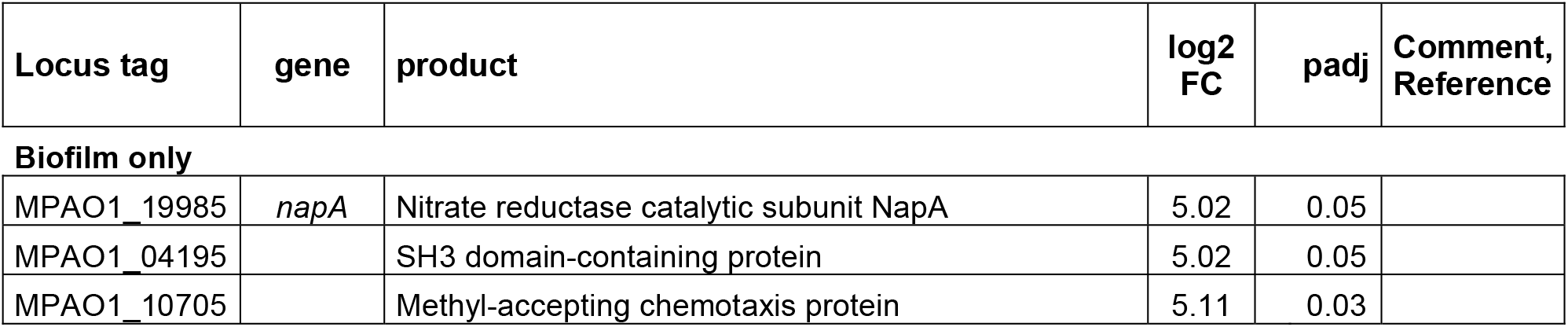
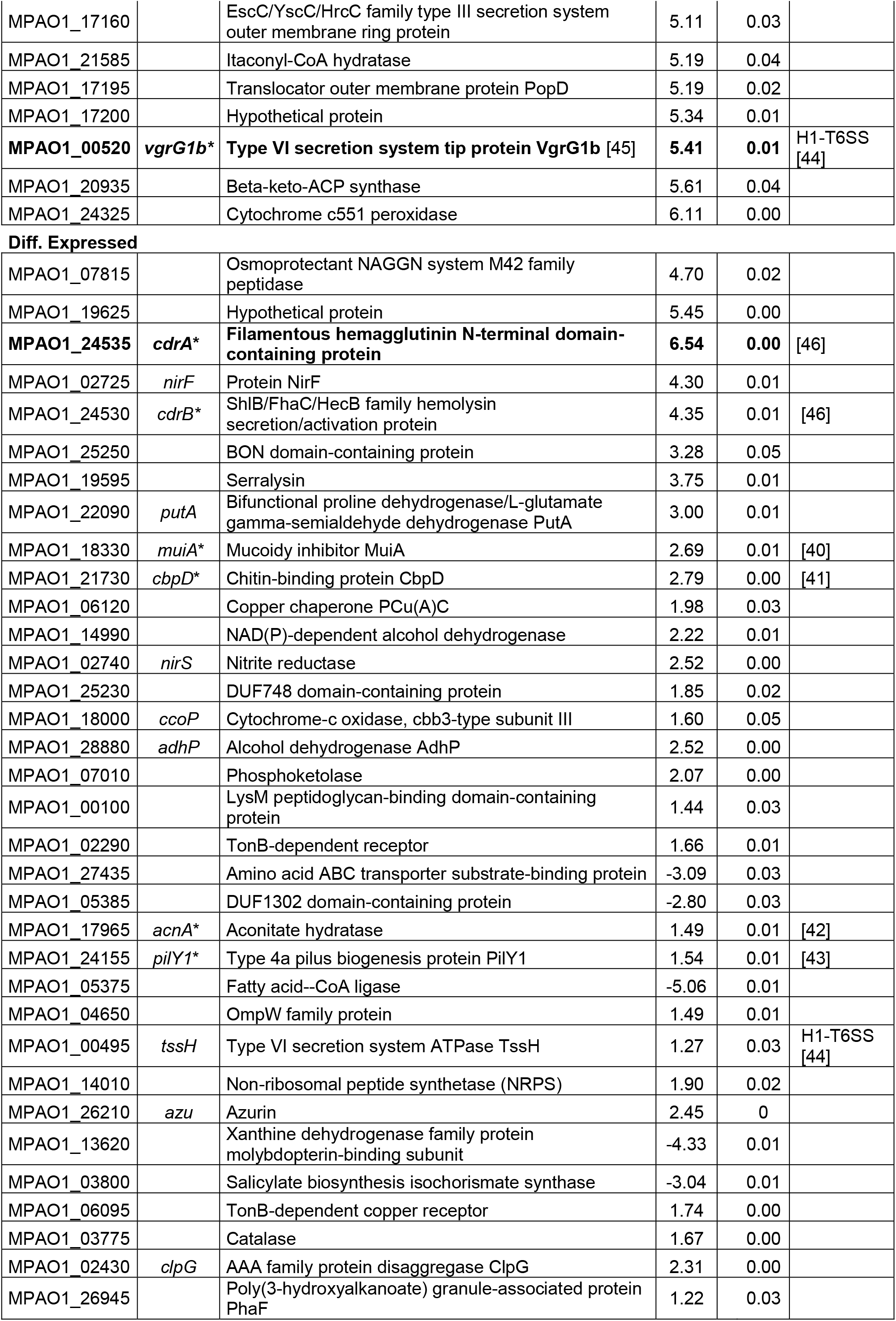
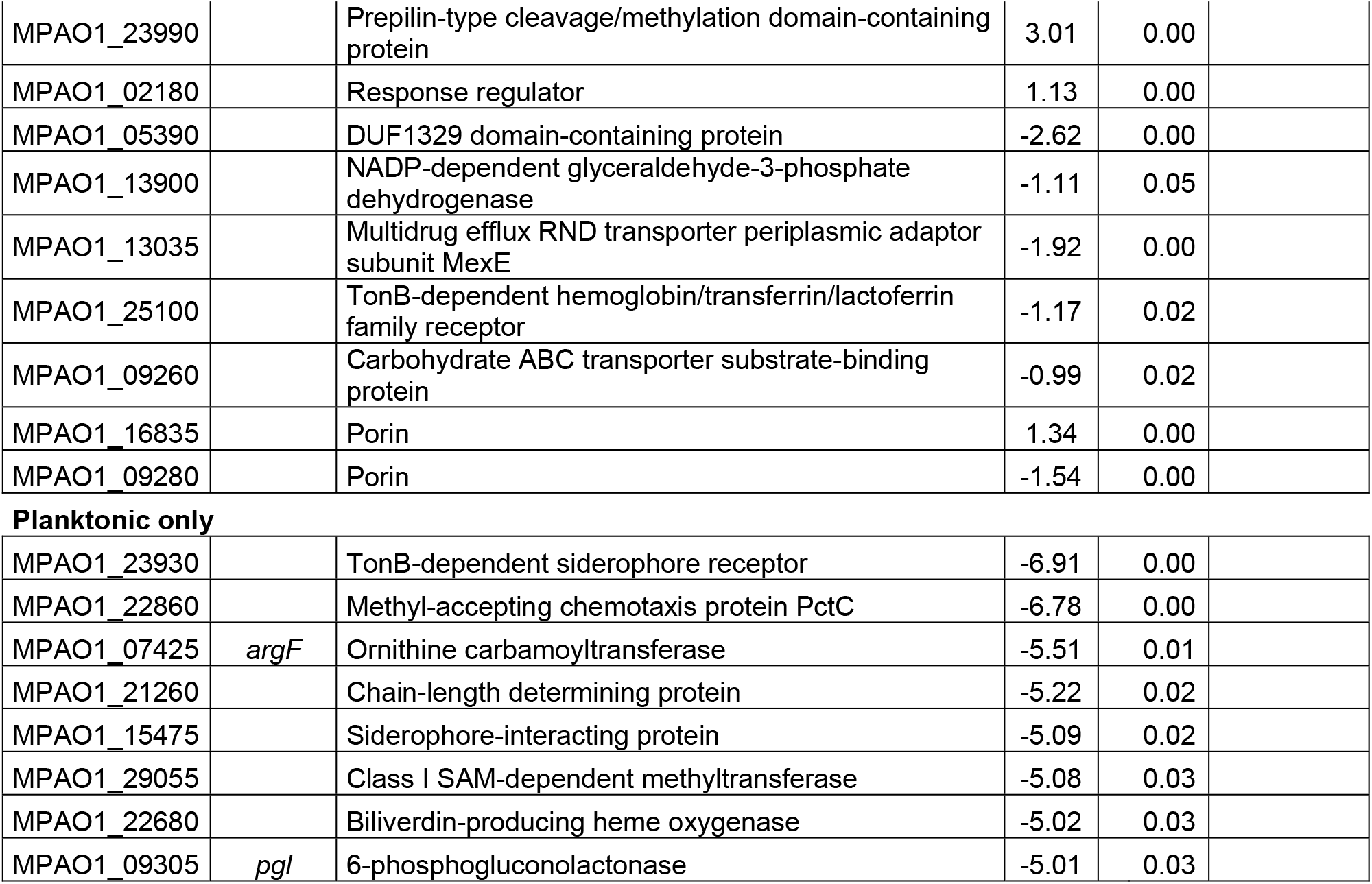
List of 61 proteins with significant differential (see text) or unique expression when comparing biofilm grown and planktonic cells. Publications linking the genes/proteins with various roles in biofilms are listed for proteins highlighted in **Fig. 5**. Two genes missed in MPAO1/P1 are shown in bold. Gene names stem from the National Center for Biotechnology Information (NCBI) annotation, or were deduced from the eggNOG annotation or the respective PAO1 homolog (*); see also Supplementary **Table 2**.

| Locus tag | gene | product | log2 FC | padj | Comment, Reference |
| --- | --- | --- | --- | --- | --- |
| **Biofilm only** |  |  |  |  |  |
| MPAO1_19985 | *napA* | Nitrate reductase catalytic subunit NapA | 5.02 | 0.05 |  |
| MPAO1_04195 |  | SH3 domain-containing protein | 5.02 | 0.05 |  |
| MPAO1_10705 |  | Methyl-accepting chemotaxis protein | 5.11 | 0.03 |  |
| MPAO1_17160 |  | EscC/YscC/HrcC family type III secretion system outer membrane ring protein | 5.11 | 0.03 |  |
| MPAO1_21585 |  | Itaconyl-CoA hydratase | 5.19 | 0.04 |  |
| MPAO1_17195 |  | Translocator outer membrane protein PopD | 5.19 | 0.02 |  |
| MPAO1_17200 |  | Hypothetical protein | 5.34 | 0.01 |  |
| **MPAO1_00520** | ***vgrG1b*\*** | **Type VI secretion system tip protein VgrG1b [45]** | **5.41** | **0.01** | H1-T6SS [44] |
| MPAO1_20935 |  | Beta-keto-ACP synthase | 5.61 | 0.04 |  |
| MPAO1_24325 |  | Cytochrome c551 peroxidase | 6.11 | 0.00 |  |
| **Diff. Expressed** |  |  |  |  |  |
| MPAO1_07815 |  | Osmoprotectant NAGGN system M42 family peptidase | 4.70 | 0.02 |  |
| MPAO1_19625 |  | Hypothetical protein | 5.45 | 0.00 |  |
| **MPAO1_24535** | ***cdrA*\*** | **Filamentous hemagglutinin N-terminal domain-containing protein** | **6.54** | **0.00** | [46] |
| MPAO1_02725 | *nirF* | Protein NirF | 4.30 | 0.01 |  |
| MPAO1_24530 | *cdrB* * | ShlB/FhaC/HecB family hemolysin secretion/activation protein | 4.35 | 0.01 | [46] |
| MPAO1_25250 |  | BON domain-containing protein | 3.28 | 0.05 |  |
| MPAO1_19595 |  | Serralysin | 3.75 | 0.01 |  |
| MPAO1_22090 | *putA* | Bifunctional proline dehydrogenase/L-glutamate gamma-semialdehyde dehydrogenase PutA | 3.00 | 0.01 |  |
| MPAO1_18330 | *muiA* * | Mucoidy inhibitor MuiA | 2.69 | 0.01 | [40] |
| MPAO1_21730 | *cbpD* * | Chitin-binding protein CbpD | 2.79 | 0.00 | [41] |
| MPAO1_06120 |  | Copper chaperone PCu(A)C | 1.98 | 0.03 |  |
| MPAO1_14990 |  | NAD(P)-dependent alcohol dehydrogenase | 2.22 | 0.01 |  |
| MPAO1_02740 | *nirS* | Nitrite reductase | 2.52 | 0.00 |  |
| MPAO1_25230 |  | DUF748 domain-containing protein | 1.85 | 0.02 |  |
| MPAO1_18000 | *ccoP* | Cytochrome-c oxidase, cbb3-type subunit III | 1.60 | 0.05 |  |
| MPAO1_28880 | *adhP* | Alcohol dehydrogenase AdhP | 2.52 | 0.00 |  |
| MPAO1_07010 |  | Phosphoketolase | 2.07 | 0.00 |  |
| MPAO1_00100 |  | LysM peptidoglycan-binding domain-containing protein | 1.44 | 0.03 |  |
| MPAO1_02290 |  | TonB-dependent receptor | 1.66 | 0.01 |  |
| MPAO1_27435 |  | Amino acid ABC transporter substrate-binding protein | -3.09 | 0.03 |  |
| MPAO1_05385 |  | DUF1302 domain-containing protein | -2.80 | 0.03 |  |
| MPAO1_17965 | *acnA* * | Aconitate hydratase | 1.49 | 0.01 | [42] |
| MPAO1_24155 | *pilY1* * | Type 4a pilus biogenesis protein PilY1 | 1.54 | 0.01 | [43] |
| MPAO1_05375 |  | Fatty acid--CoA ligase | -5.06 | 0.01 |  |
| MPAO1_04650 |  | OmpW family protein | 1.49 | 0.01 |  |
| MPAO1_00495 | *tssH* | Type VI secretion system ATPase TssH | 1.27 | 0.03 | H1-T6SS [44] |
| MPAO1_14010 |  | Non-ribosomal peptide synthetase (NRPS) | 1.90 | 0.02 |  |
| MPAO1_26210 | *azu* | Azurin | 2.45 | 0 |  |
| MPAO1_13620 |  | Xanthine dehydrogenase family protein molybdopterin-binding subunit | -4.33 | 0.01 |  |
| MPAO1_03800 |  | Salicylate biosynthesis isochorismate synthase | -3.04 | 0.01 |  |
| MPAO1_06095 |  | TonB-dependent copper receptor | 1.74 | 0.00 |  |
| MPAO1_03775 |  | Catalase | 1.67 | 0.00 |  |
| MPAO1_02430 | *clpG* | AAA family protein disaggregase ClpG | 2.31 | 0.00 |  |
| MPAO1_26945 |  | Poly(3-hydroxyalkanoate) granule-associated protein PhaF | 1.22 | 0.03 |  |
| MPAO1_23990 |  | Prepilin-type cleavage/methylation domain-containing protein | 3.01 | 0.00 |  |
| MPAO1_02180 |  | Response regulator | 1.13 | 0.00 |  |
| MPAO1_05390 |  | DUF1329 domain-containing protein | -2.62 | 0.00 |  |
| MPAO1_13900 |  | NADP-dependent glyceraldehyde-3-phosphate dehydrogenase | -1.11 | 0.05 |  |
| MPAO1_13035 |  | Multidrug efflux RND transporter periplasmic adaptor subunit MexE | -1.92 | 0.00 |  |
| MPAO1_25100 |  | TonB-dependent hemoglobin/transferrin/lactoferrin family receptor | -1.17 | 0.02 |  |
| MPAO1_09260 |  | Carbohydrate ABC transporter substrate-binding protein | -0.99 | 0.02 |  |
| MPAO1_16835 |  | Porin | 1.34 | 0.00 |  |
| MPAO1_09280 |  | Porin | -1.54 | 0.00 |  |

Planktonic only
|  |  |  |  |  |
| --- | --- | --- | --- | --- |
| MPAO1_23930 |  | TonB-dependent siderophore receptor | -6.91 | 0.00 |
| MPAO1_22860 |  | Methyl-accepting chemotaxis protein PctC | -6.78 | 0.00 |
| MPAO1_07425 | *argF* | Ornithine carbamoyltransferase | -5.51 | 0.01 |
| MPAO1_21260 |  | Chain-length determining protein | -5.22 | 0.02 |
| MPAO1_15475 |  | Siderophore-interacting protein | -5.09 | 0.02 |
| MPAO1_29055 |  | Class I SAM-dependent methyltransferase | -5.08 | 0.03 |
| MPAO1_22680 |  | Biliverdin-producing heme oxygenase | -5.02 | 0.03 |
| MPAO1_09305 | *pgl* | 6-phosphogluconolactonase | -5.01 | 0.03 |

Notably, while the overall number of predicted genes was close for both strains (**Table 1**), we observed 232 gene clusters specific to strain MPAO1 and 21 clusters specific to strain PAO1-UW (**Fig. 1c**), suggestive of potentially relevant differences between the strains. The annotation of the shared (core) and strain-specific (unique) gene clusters is provided in Supplementary **Table 4**. This analysis indicated that a sizeable set of genes were specific to the MPAO1 genome, and that mapping datasets obtained from this strain back to the PAO1-UW genome could overlook important genes (see below). A gene ontology (GO) enrichment analysis of the MPAO1 unique proteins against all CDS in its genome revealed that the biological process “protein phosphorylation” was significantly enriched (*p* value < 0.01) with 10 hits among all genes including three among the unique genes (including a DNA helicase and 2 serine/threonine protein kinases; Supplementary **Table 5**). Furthermore, for the biological process “Bacteriocin immunity” five hits were found among all genes, two of which were among the unique MPAO1 genes (Supplementary **Table 5**).

### Tn-seq data mapping

The complete MPAO1 genome sequence allowed us to re-analyze public Tn-seq datasets without the limitation of any remaining “genomic blind spots” that otherwise might preclude an identification of all essential genes [25], and the drawbacks of mapping Tn-seq data to a closely related reference genome. A re-mapping of MPAO1 Tn-seq datasets obtained from several conditions (LB medium, minimal medium, sputum and brain-heart infusion BHI medium) [24] against both the PAO1-UW genome and our MPAO1 genome (see Methods), confirmed our expectation. We indeed observed a higher percentage of mapped reads for MPAO1 (roughly 0.1 - 0.35% of all mapped reads per sample; Supplementary **Table 6**) and unique insertion sites (roughly 0.2% more in MPAO1, Supplementary **Table 6**). Genes with no insertion or genes whose *p* value was less than 0.001 were considered essential (see Methods). Overall, 577 genes were classified as essential in one of the three primary growth conditions LB medium, minimal medium, sputum (Supplementary **Table 7**), and 312 genes represented general essential genes, i.e., were essential in all three growth conditions, respectively (Supplementary **Fig. 2**). Importantly, close to 40 MPAO-1 unique genes were linked here for the first time with an essentiality status, as they were essential in one or more of the 16 Tn-seq libraries (Supplementary **Table 7**). By mapping data against the PAO1-UW genome, these genes had been previously overlooked in the analysis of essential *P. aeruginosa* genes.

**Fig. 2.**
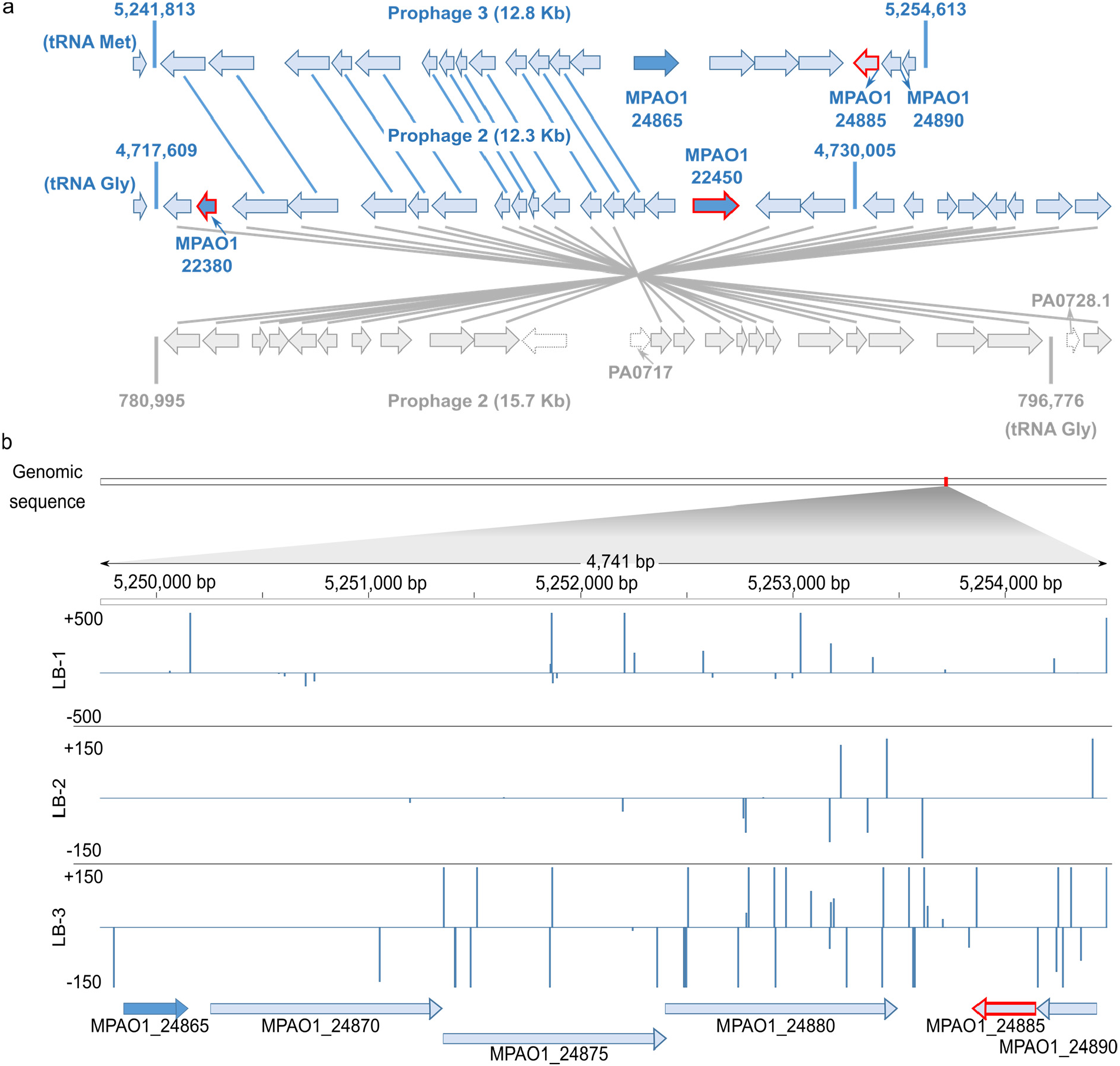
An overview of annotated genes in selected prophage regions and their essentiality classification. MPAO1-unique essential genes are shown in dark blue, general essential MPAO1 genes with a red arrow outline. (**a**) Genes located in prophage region 2 of PAO1-UW (gray), the corresponding inverted region in strain MPAO1 (light blue arrows in middle), and the prophage region 3 (light blue arrows on top) unique to MPAO1 are shown (not drawn to scale), the genomic positions of their boundaries (5’ to 3’) and flanking tRNAs. Genes connected by lines are orthologous to each other based on comparative genomics combined with a Blast analysis. (**b**) Transposon insertions in selected genes of prophage region 3 of MPAO1. Insertion frequencies in six genes are shown using data mapped from the LB-1 (3 replicates), LB-2 (2 replicates) and LB-3 (1 sample) Tn-seq libraries. Non-essential genes (based on dataset of 577 genes essential in one of three primary growth conditions) are shown in light blue.

Among these MPAO1-unique genes, we identified 18 genes that were essential in 50% or more of the Tn-seq runs, six of which represented general essential genes (**Table 2**). The general essential genes included two genes located in the prophage2 region, i.e., MPAO1_22380, a type II Phd/YefM family antitoxin gene located next to MPAO1_22375, coding for a RelE/ParE type toxin, and MPAO1_22450, a DNA-binding protein (**Fig. 2a**; arrows framed in red). A further general essential gene was MPAO1_00215 encoding for a hypothetical protein. MPAO1_00215 is located in a genomic region that harbors another essential gene (MPAO1_00230, Supplementary **Table 2**), that may represent an operon.

Furthermore, the prophage 3 region unique to strain MPAO1, harbored a gene encoding a hypothetical protein (MPAO1_24865; **Fig. 2b**) that was essential in eight of 16 samples (Table 2). Conversely, MPAO1_24885 (addiction module antidote protein from the HigA family toxin-antitoxin (TA) system) from this region was even classified as general essential (**Table 2**; 14 of 16 samples); due to its homology to PA4726, it is not unique to MPAO1. (**Fig. 2b**). Together with the non-essential MPAO1_24890 (plasmid maintenance system killer protein; most similar to RelE-like toxins of the type II TA system HigB), MPAO1_24885 encodes for a TA system. The addiction module protein MPAO1_24885 is homologous to PA4674 in PAO1-UW, which is among the list of 352 general essential genes reported by Lee and colleagues and encodes the HigA antitoxin [24]. However, there is no homolog annotated for the plasmid maintenance killer protein MPAO1_24890 in PAO1-UW. Therefore, due to this missing gene, the TA system was not identified in PAO1-UW. This finding again underlines the importance of having the actual and complete genome sequence to map functional data.

### Reproducible formation of MPAO1 biofilms

The second important objective of our integrated model system was to enable the reliable generation of phenotypic data under biofilm-growth relevant conditions. For this purpose, we focused on the development of a microfluidic flow chamber for reproducible biofilm formation that would allow us to subsequently identify genes relevant for biofilm growth and AMR development. The flow chamber was designed in such a way that we could assess the effects of hydrodynamic conditions [35], such as shear stress and controlled flow conditions. Our flow chamber was replicated in PDMS, a simple to use, transparent and breathable elastomer material that naturally adheres to glass. A straight microfluidic channel design was used (30 mm length x 2 mm width x 0.200 mm depth) (**Fig. 3a**, see Methods for further details). PDMS was selected due to its broad application in indwelling devices and implant materials [36]. The inlet and outlet of the microfluidic flow chamber comprised of sterile polytetrafluoroethylene (PTFE) tubing, a material that was chosen because it generally exhibits low bacterial adhesion. A syringe pump was used to deliver 5 μL/min (*u*≈200 mm/s) flow inside the chamber to provide laminar flow conditions for bacterial adhesion and biofilm growth (the calculated Reynolds Number corrected for transport of water at 37°C was 0.103; for details see Supplementary **Table 8**).

**Fig. 3.**
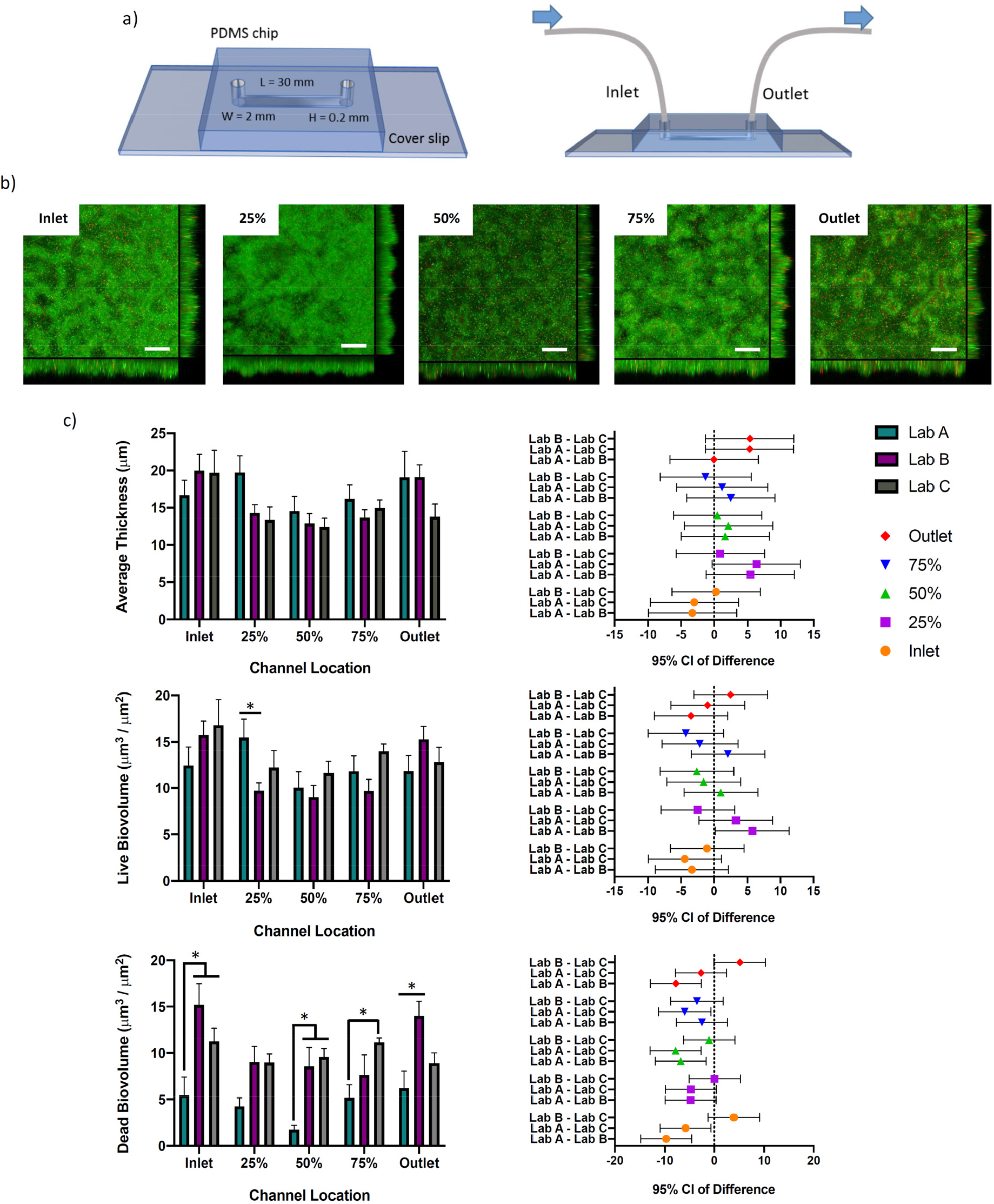
The publicly available mold design for the microfluidic flow chamber allows reproducible biofilm formation as confirmed by an inter-laboratory comparison. (**a**) Schematic and dimensions of the flow chamber. (**b**) Representative images of 72 h MPAO1 WT biofilms grown on the PDMS surface of the device under laminar flow conditions at five different locations along the channel. Biofilms were treated with live/dead staining (green – live cells stained with Syto9; red – dead cells stained with propidium iodide). Scale bar in confocal XY plane: 40 µm. Sagittal XZ section represents biofilm thickness. (**c**) COMSTAT data for average thickness, and live/dead biovolume of 72 h MPAO1 WT biofilms generated by three different laboratories, with 95% confidence interval comparisons (3 biological repeats comprising 3 technical repeats per site, i.e., n=9 biological / n=27 technical repeats overall; error bars -standard error of mean; 2-way ANOVA with lab and channel location as variables followed by multiple comparisons Tukey test). \**p* value < 0.05.

The reproducibility of a 72 h mature MPAO1 biofilm on the PDMS surface of the device was investigated by confocal laser scanning microscopy (CLSM) combined with live/dead staining using the dyes Syto9 and propidium iodide in three separate consortium laboratories all using the same microfluidic chamber mold (design publicly available; see Data Access) **(Fig. 3b, c)**. The biofilms formed in the three laboratories were consistent with data falling within 95% confidence intervals, the only difference being the observation of a reduced dead biovolume in one laboratory’s model (Lab A; *p* value < 0.05). Biofilm formation was relatively uniform throughout the flow channel with an average thickness of 16 µm and a small reduction observed towards the center of the channel (Inlet - 18.8 µm, 25% - 15.8 µm, 50% - 13.3 µm, 75% - 14.9 µm, Outlet - 17.3 µm). An average biovolume of 12.5 µm^3^/µm^2^ and dead biovolume of 8.4 µm^3^/μm^2^ was observed, again reducing towards the center of the device commensurate with the average biomass.

### Screening experiments identify known and new genes relevant for biofilm formation and antibiotic resistance

The MPAO1 transposon mutant library was tested with a 96-well plate screening system that was devised to enable the identification of genes that affect biofilm formation and/or play a role in the development of AMR. A batch of fifty randomly selected mutants (see Supplementary **Table 9)** was taken from the library to first test the reliability of our protocol to identify genes related to biofilm formation (in duplicate). Strain PW8965 harboring an insertion in *cbrB* (PAO1 identifier PA4726, MPAO1_25185), a transcriptional activator that forms part of the CbrA/CbrB two-component system important in catabolite repression [37], was found to produce the least amount of biofilm (**Fig. 4a**). Three independent experiments (**Fig. 4a**, right panel) confirmed that the *cbrB* mutant produced significantly less biofilm biomass (*p* value < 0.001) compared to the WT and strain PW7021 (an *arnB* mutant; see below). Biofilm growth of the *cbrB* mutant was also performed within the flow chamber to confirm the capacity of the device to assess differential biofilm formation. Similar to the 96-well plate screening assay, the *cbrB* mutant produced substantially less biofilm compared to the MPAO1 WT over 18 h in the flow chamber **(Fig. 4c)**.

**Fig. 4.**
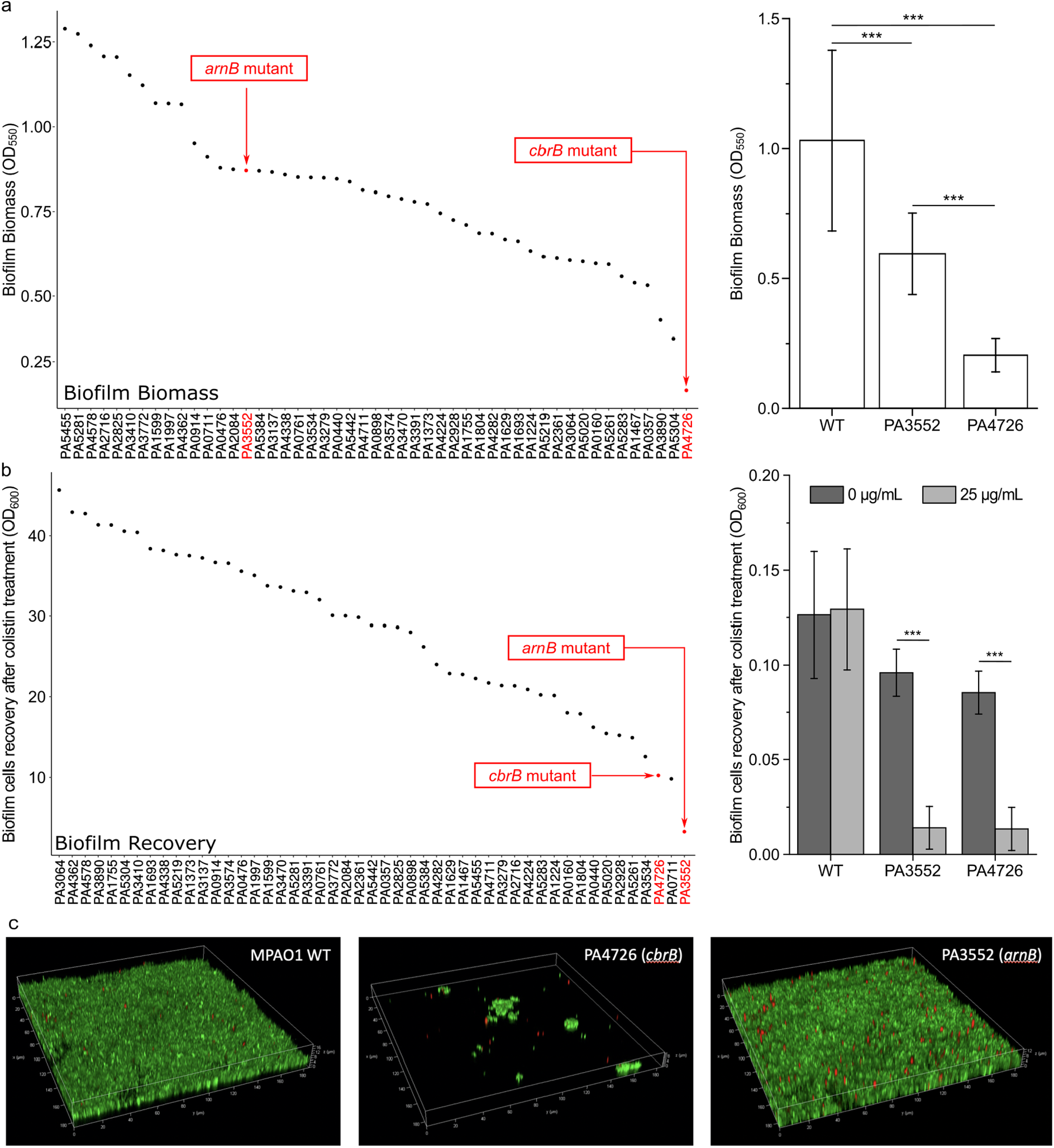
Proof of principle that biofilm growth-relevant and AMR-related genes can be identified in adequate screens using the MPAO1 transposon mutant library. (**a**) Biofilm formation of fifty MPAO1 mutant strains (X-axis) after 24h incubation in M9 medium (average of two independent wells). Biofilm biomass was quantified by crystal violet. The *cbrB mutant* demonstrated substantially reduced biofilm formation in the screen compared to relatively robust biofilm formation for the *arnB mutant*. The right panel shows a statistic evaluation of three replicates from WT, *cbrB* and *arnB* cells (***; *p* value < 0.001). (**b**) Ability of biofilms formed by fifty MPAO1 mutant strains to recover after colistin treatment (see Methods). The recovery of treated biofilm cells was normalized to the recovery of non-treated biofilm cells (defined as 100%). Both *cbrB* and *arnB mutants* demonstrated poor recovery after colistin treatment. Analysis of three replicates uncovered statistically significant differences (***; *p* value < 0.001). (**c**) Comparative confocal micrographs after live/dead staining (green – live cells stained with Syto9; red – dead cells stained with propidium iodide) of 18 h MPAO1 WT, *cbrB* and *arnB* biofilms grown under microfluidic conditions confirm reduced biofilm formation for the *cbrB mutant* and robust biofilm formation of the *arnB mutant* in the absence of treatment.

Next, we tested the fifty strains for their biofilm resistance to colistin. Strain PW7021, harboring an insertion in *arnB* (PA3552; MPAO1_07345) was included as positive control. ArnB is a well-studied protein known to modify lipopolysaccharide (LPS) and play a key role in the resistance to colistin [38, 39]. The recovery of biofilm cells after treatment with 25 µg/ml colistin was compared to the recovery of non-treated biofilm cells (**Fig. 4b**) (see Methods), as described previously [13]. This concentration of colistin was much higher than the minimal inhibitory concentration (MIC) used for the planktonic *P. aeruginosa* MPAO1 (4 µg/mL) allowing us to focus specifically on the biofilm cells. As expected, the *arnB* mutant exhibited a very low recovery after colistin treatment (97% less than the control without colistin) (**Fig. 4b**). In contrast, the *arnB* mutant produced robust biofilms in the biofilm screening assay (**Fig. 4a**), a phenotype that was confirmed using the microfluidic chamber (**Fig. 4c**). Notably, the *cbrB* mutant strain grown as a biofilm was also found to be sensitive to colistin (90% less recovery than the control without colistin; **Fig. 4a**), which might be related with the low amount of biofilm produced by this mutant. An independent repetition in triplicate confirmed the significant sensitivity of the *arnB* and *cbrB* mutants compared to the WT (**Fig. 4b**, right panel) and showed that biofilm cells of these two strains are killed with doses down to 12.5 µg/mL of colistin (data not shown).

### Protein expression profiling of MPAO1 grown planktonic and in biofilms

To assess if we could identify proteins known to play a role in biofilm formation with the microfluidic chamber, we next generated shotgun proteomics data for MPAO1 cells grown to mid-exponential planktonic phase or as 72 h biofilms (3 replicates each). 1,530 and 1,728 proteins were identified in planktonic cells and biofilm, respectively, resulting in a combined 1,922 of the 5,799 annotated proteins (33.1%). Among the most significantly differentially expressed proteins (log_2_ fold change (FC) of ≥ 1 or ≤ -1 and adjusted *p* value ≤ 0.05; see Methods) several candidates were identified that have previously been linked with a role in biofilm formation. These included MuiA (MPAO1_18330) [40], CbpD (MPAO1_21730) [41], AcnA (MPAO1_17965) [42] and PilY1 (MPAO1_24155) [43] (**Fig. 5a, Table 3**; see Discussion). In addition, the hypothetical protein MPAO1_19625 was highly upregulated in biofilms (**Fig. 5a**), indicating that hypothetical proteins can be linked to roles in biofilm formation and growth. We next looked for proteogenomic evidence of genomic differences and found that 21 of the 52 CDSs that were missed in the fragmented MPAO1/P1 assembly were expressed at the protein level (Supplementary **Table 2**). Notably, this included two proteins significantly upregulated in the biofilm, namely MPAO1_00520 (T6SS tip protein VgrG1b) located close to the H1 type VI secretion system (T6SS) cluster [44] and MPAO1_24535 (**Fig. 5a**), the homolog of PAO1 CdrA, a cylic-di-GMP-regulated adhesin known to reinforce the biofilm matrix [46], again underlining the importance of a complete genome sequence for downstream functional genomics analyses. Notably, nine of 14 structural genes of H1-T6SS, one of overall three T6SSs in *P. aeruginosa* that helps it to prevail in challenging niches [47], were upregulated around two-fold or more in biofilm (Supplementary **Fig. 3**). Similarly, all three VgrG1 proteins (1a-1c) that are co-regulated with the H1-TS66 [45] were upregulated in biofilm, while none of the other seven additional VgrG family members was expressed.

**Fig. 5.**
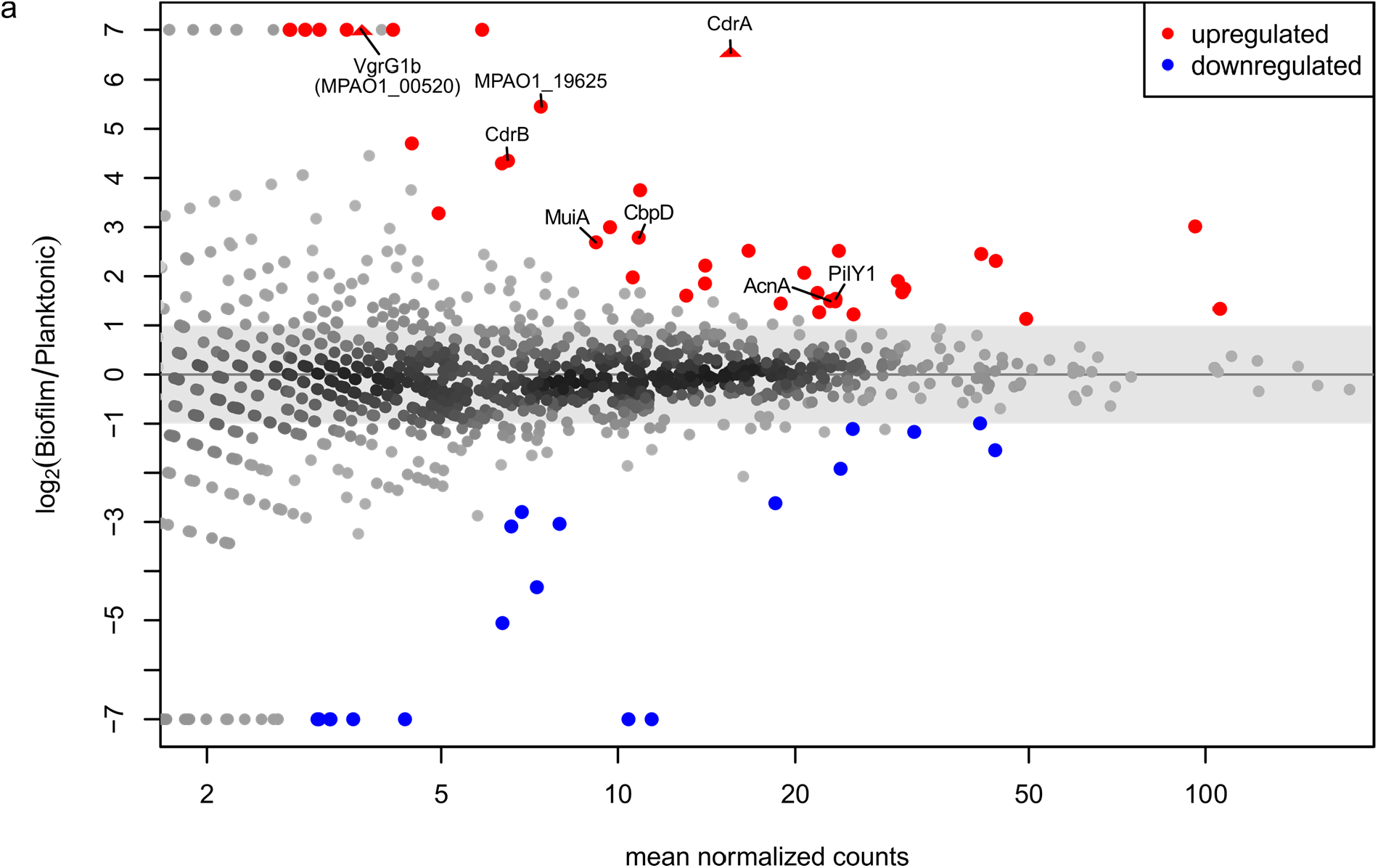
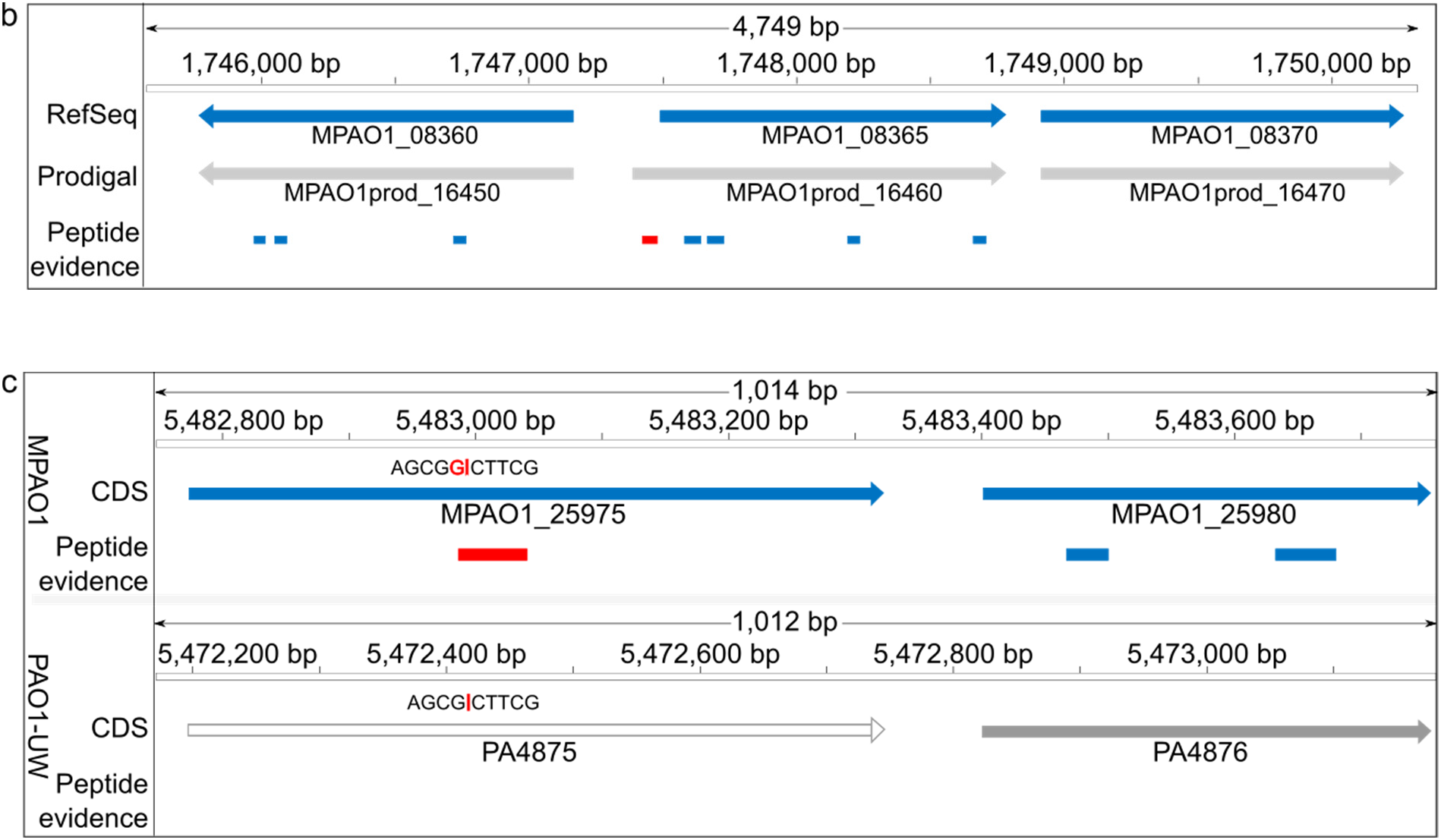
Proteomic experiments identify known biofilm-related proteins and novel information. (**a**) Differential protein expression between MPAO1 mid-exponential planktonic cells and 72 h biofilms. Selected significantly upregulated proteins (red dots) known to play a role in biofilm formation/growth are labeled, proteins downregulated in planktonic growth are shown in blue. Red triangles denote proteins encoded by genes missed in the MPAO1/P1 genome. (**b**) Proteogenomic expression evidence for a longer protein than annotated by RefSeq: the Prodigal predicted protein MPAO1prod_16460 (gray arrow; 447 aa; amino acid) is 44 aa longer than the RefSeq annotated MPAO1_08365 and encodes a glutamine synthetase (blue arrow; 413 aa). The NH-terminal extension is supported by 1 peptide (red) with seven PSMs and harbors a 40 aa longer glutamine synthetase N-terminal domain compared to the RefSeq protein. (**c**) Proteogenomic expression evidence for a single nucleotide insertion (red) in the MPAO1_25975 gene (blue arrow) compared to its PAO1 homolog PA4875 (annotated as pseudogene; gray open arrow). The change is supported by peptide evidence (1 red bar).

Finally, to identify unannotated short ORFs that may carry out important functions or novel start sites by proteogenomics, we created an integrated proteogenomics search database (iPtgxDB) for strain MPAO1(Supplementary **Table 10**), which covers its entire coding potential [31]. A search combined with stringent result filtering (see Methods) allowed us to identify unambiguous peptide evidence [48] for a 44 aa longer proteoform of MPAO1_08365 (predicted by Prodigal, an *ab initio* gene prediction algorithm; **Fig. 5b**), as well as proteogenomic evidence supporting an SNP in strain MPAO1 (**Fig. 5c**). Compared to PA4875 (annotated as pseudogene in strain PAO1), the corresponding MPAO1 homolog (MPAO1_25975) harbored a single nucleotide insertion. The peptide that supported this single nucleotide change at the amino acid level was identified with seven peptide spectrum matches (PSMs), illustrating the ability to identify SNP changes at the protein level, with implications for clinical proteomics.

## Discussion

*P. aeruginosa* is a member of the ESKAPE pathogens, the lead cause of worldwide nosocomial infections [10]. Along with many other clinically relevant bacteria, it can form biofilms whose emergent properties [49] include a much higher tolerance to antimicrobials. Together with the increased mutation rates in biofilm compared to planktonic cells [17], this further complicates treatment and cure of biofilm-based infections [12, 13]. The development of model systems allowing the study of antimicrobial tolerance mechanisms and the evolutionary dynamics that lead to AMR development in biofilms is thus of utmost priority.

We here develop and validate such a model system for *P. aeruginosa* MPAO1 (**Fig. 6**). Conceptually, the model was designed to integrate genotype data with phenotypic information and to leverage the wealth of existing public genetic resources and functional genomics datasets. A complete, fully resolved genome sequence is one critical element [31, 50]. While this existed for *P. aeruginosa* PAO1 [2], only three fragmented Illumina-based genome assemblies were available for MPAO1, the parental strain of the popular UW transposon mutant library [21]. These included strains MPAO1/P1 [32] and the recently sequenced PAO1-2017-E and PAO1-2017-I [19]. On average, they lacked between 55 to 66 genes (40 to 52 CDS) compared to our complete MPAO1 genome (Supplementary **Table S2**). For MPAO1/P1, these included the essential *ftsY*, an adhesin and several T6SS effectors (see below), and four of the overall eight NRPSs. NRPSs are highly relevant for AMR as they often represent enzymes involved in the biosynthesis of antibiotics [51]. In fact, due to the multi-resistant phenotype of ESKAPE pathogens, concerted efforts aim to describe their NRPS gene clusters in search for novel therapeutic approaches [52].

**Fig. 6.**
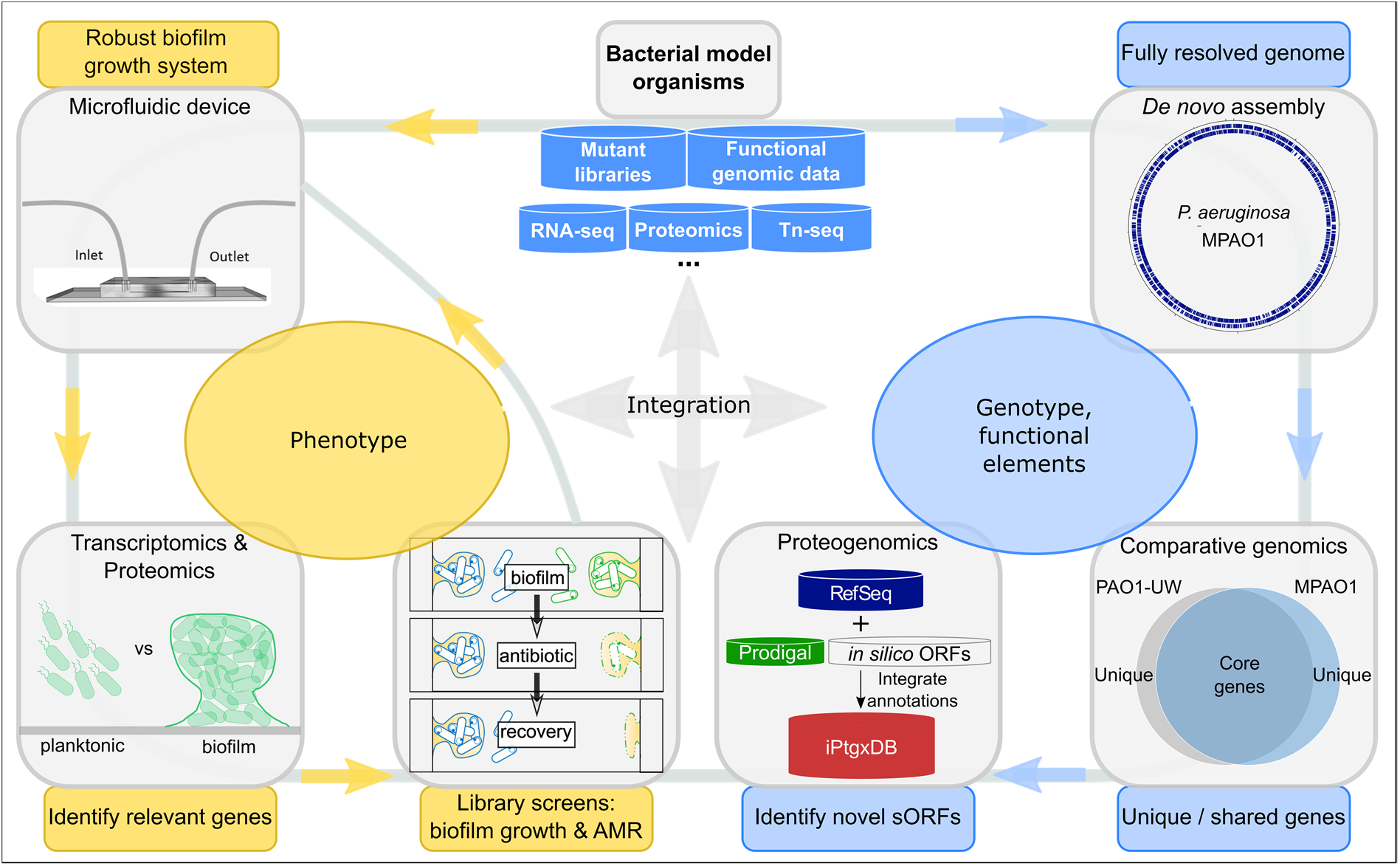
Integrated model system to identify and validate genes relevant for biofilm growth and AMR. A sequential genomics-driven workflow (blue arrows) to *de novo* assemble the complete genome, identify unique and conserved genes among key reference strains by comparative genomics and missed genes by proteogenomics is integrated with an experimental workflow in the form of an iterative cycle that can be entered at various points (yellow arrows). This workflow allows the study of biofilm grown cells, to explore differentially expressed genes or proteins compared to planktonic cells and to screen mutant libraries to identify functionally relevant genes. The model leverages the enormous value of genetic resources like gene knockout or transposon insertion mutant libraries and functional genomics datasets (RNA-seq, Tn-seq, etc.; blue containers). Additionally, it allows for phenotypic characterization of biofilms formed by mutant strains, thereby allowing us to determine the impact of specific genes on biofilm formation and assess their role in AMR (yellow arrows).

Comparative genomics with the PAO1 type strain uncovered an inventory of conserved and strain-specific genes, and a list of genome-wide SNPs, extending an earlier study that had compared a subset of genomic regions [20]. Among the 232 MPAO1-unique gene clusters, bacteriocins [53] were enriched, which play a role in restricting the growth of closely related microbial competitors to gain an advantage in colonizing a variety of niches [54]. The complete MPAO1 genome enabled us to remap valuable existing Tn-seq datasets from relevant conditions [24], thereby identifying 39 MPAO1 essential genes that had escaped detection so far due to reference-based PAO1 mapping. 18 thereof were essential in at least 50% of the16 Tn-seq samples, and six represented general essential genes, including a Phd/YefM family type antitoxin (MPAO1_22380), which was essential in all samples. Importantly, our data do not conflict with results from previous studies; rather, they open the field to study the roles of additional MPAO1-unique essential genes. Furthermore, our results suggest that groups planning to construct inventories of core essential genes in other pathogens, following the elegant approach of Poulsen et al. [25], should ideally select complete genomes without any genomic blind spots. They had considered both relevant media mimicking different infection types and nine strains from different lineages of a *P. aeruginosa* phylogenetic tree, to identify 320 core general essential genes as a high priority drug target list [25].

To leverage the experimental arm of our model (**Fig. 6**), the consortium first developed a PDMS microfluidic flow chamber for biofilm growth, which offers several significant advantages. It provides laminar flow conditions inside the channels (Supplementary **Table 8**), allows gas exchange, decreases the amount of growth medium, facilitates heat transfer, is inexpensive to replicate and permits imaging of the biofilm and easy harvesting for biochemical characterization. While the flow chamber can be used to monitor biofilm formation on both glass (oxygen impermeable) and PDMS, it is more relevant to investigate biofilm formation on PDMS as a widely applied biomaterial used in indwelling devices and implants [36]. The fact that PDMS can transport oxygen to the base of the biofilm can potentially lead to different biofilm characteristics; we observed that biofilms on PDMS formed a more homogeneous layer (**Fig. 3b**) as compared to the commonly observed mushroom-like structures of *P. aeruginosa* biofilms [55]. With an increasing number of coatings being developed for future use in the clinics [56], we plan to exploit the advantage of PDMS in that different antimicrobial compounds can easily be mixed into the polymer before curing, providing a simple model to study the effect of different antimicrobial coatings on AMR in a defined biofilm model.

The microfluidic data from the interlaboratory trial on strain MPAO1 validated the utility of the flow chamber and allowed us to compare the phenotypes of WT and mutant strains of the UW transposon library. Important genes were first identified with a microtiter plate screening assay and subsequently validated with the flow chamber. Proof of principle experiments confirmed the role of *arnB* (PA3552), i.e., a gene relevant for colistin resistance [38, 39], both in biofilms grown in the 96-well plate screen and the flow chamber. In addition, a mutant lacking *cbrB* (PA4726) showed reduced resistance to colistin in biofilm and formed very low amounts of biofilm in both the microtiter plate and flow chamber. As part of the two-component system CbrAB, a mutation in the response regulator *cbrB* is known to negatively affect the use of several carbon and nitrogen sources [37]. Such a defect could explain the low growth rate (data not shown), the low biofilm biomass and therefore the low resistance to colistin of this mutant. Using *P. aeruginosa* PA14, it was shown that a mutation in CbrA improved biofilm formation, while a mutation in CbrB did not [57]. However, these differences might be explained by strains (MPAO1 versus PA14) or growth media used (M9 versus BM2-biofilm medium). Together, the screens demonstrated that known and novel genes related to AMR and biofilm formation can be identified and validated.

The differential proteomics data confirmed proteins known to play a role in biofilm formation and growth. These included MuiA, which inhibited swarming motility and enhanced biofilm formation (roles, that were validated in knockout strains) [40], and CbpD, for which higher protein expression had been observed in late phases of biofilm growth; accordingly, mutants displayed a lower amount of biofilm growth and exopolysaccharides (EPS) [41]. Inactivation studies showed that the gene encoding AcnA impaired biofilm formation and was required for microcolony formation [42], while increased expression of PilY1 repressed swarming and increased biofilm formation, as confirmed by knockout experiments [43]. Biofilm exclusive protein expression was observed for MPAO1_00520, the T6SS VgrG1b effector protein [45], while the adhesin CdrA (MPAO1_24535) [46] was highly upregulated in biofilms. Both genes were missed in the MPAO1/P1 genome. CdrA forms a two-partner secretion system with CdrB, and both were upregulated under elevated c-di-GMP levels [46], in line with the upregulation we observed in biofilm. Moreover, an NRPS (MPAO1_14010) and the hypothetical protein MPAO1_19625 were significantly upregulated in biofilm (**Table 3**). The data provided insights beyond the top differentially expressed proteins. Notable examples included immunity protein TplEi [58] (PA1509, MPAO1_18250), a bacteriocin of the H2-T6SS [47], which was exclusively expressed in biofilm (Supplementary **Table 2**), and upregulation of nine of 14 structural members of H1-T6SS [47] (Supplementary **Fig. 3**). Active T6SSs have been associated with chronic infections in cystic fibrosis patients [45], and H1-T6SS plays an important role in dominance of *P. aeruginosa* in multi-species biofilms [59].

The public MPAO1 iPtgxDB allows to identify missed genes by proteogenomics [31], which often encode short proteins (sProteins) that can carry out important functions [60, 61]. Interestingly, Tn-seq data from the Manoil group had implied an essential genomic region in the PF1 phage region of PAO1-UW [24]. Re-mapping their data, we identified a general essential gene (MPAO1_22380) annotated in our MPAO1 genome whose homolog had been missed in the PAO1 genome annotation, and which appeared to encode the antitoxin member of a ParDE-like TA system (PA0728.1, **Fig. 2**). Unfortunately, we did not identify expression evidence for the antitoxin MPAO1_22380 (83 aa) with our iPtgxDB, most likely because our dataset (33% of MPAO1 proteins) was not as extensive as that used in a comprehensive proteogenomic study (85% of *Bartonella henselae* proteins) [31], whose complete membrane proteome coverage included expression evidence for all T4SS members [62]. Nevertheless, we observed proteogenomic evidence for gene products missed in the fragmented MPAO1/P1 genome, for new start sites and for single amino acid variations, underlining the potential value of proteogenomics for application in clinical proteomics.

Our proof of principle experiments uncovered several candidates for follow-up studies and illustrated the benefit of the complete MPAO1 genome, which lead to the discovery of six general essential genes not contained in the transposon library, and which will allow to identify evolutionary changes that lead to AMR in biofilm by deep sequencing. Having been validated for the generation of reproducible inter-laboratory *P. aeruginosa* biofilm results, a milestone en route to a community standard (see Data Access), the microfluidic platform can be instrumental to investigate other biofilms, notably clinical pathogens and mixed-species biofilms [59]. The upregulation of the H1-T6SS highly relevant for dominance of *P. aeruginosa* [59] implies that our microfluidic chamber should be valuable also for this extension. Our proposed workflow (**Fig. 6**) with feedback between genotypic and phenotypic assessment of biofilm characteristics can thus be leveraged across the field of biofilm research and helps bridge the gap between genome-wide and reductionist approaches to study the role of biofilms in AMR development.

## Methods

### Bacterial growth and genomic DNA extraction

*P. aeruginosa* strain MPAO1 (originating from the lab of Dr. Barbara Iglewski) was obtained from Prof. Colin Manoil, UW (Seattle, USA) together with the transposon insertion mutant collection of ∼5000 mutated genes [21]. For DNA extraction, the MPAO1 cryoculture was streaked out on 20% BHI solid medium (7.4 g in 1 L water) containing 1.5 % agar (both Sigma, Switzerland). Shaken 20% BHI fluid cultures were inoculated from a single colony and grown at 30 °C until mid-exponential phase (OD600 = 0.5). Genomic DNA (gDNA) was extracted with the GeneElute kit (Sigma, Switzerland), following the Gram-negative protocol, including RNase treatment. An analysis of 9331 complete bacterial genomes (February 23, 2018) indicated that 106 *P. aeruginosa* strains had been sequenced completely [29]. Only two were PAO1 strains, 38 had very difficult to assemble genomes with repeat pairs greater 10 kilo base pairs (bp).

### Sequencing, *de novo* genome assembly and annotation

PacBio SMRT sequencing was carried out on a RS II machine (1 SMRT cell, P6-C4 chemistry). A size selection step (BluePippin) was used to enrich for fragments longer than 10 kb. The PacBio run yielded 105,221 subreads (1,32 Gbp sequence data). Subreads were *de novo* assembled using the SMRT Analysis portal v5.1.0 and HGAP4 [63], and polished with Arrow. In addition, a 2 x 300 bp paired end library (Illumina Nextera XT DNA kit) was sequenced on a MiSeq. Polishing of the assembly with Illumina reads, circularization, start alignment using *dnaA* and final verification of assembly completeness were performed as described previously [64]. The quality of the aligned reads and the final chromosome was assessed using Qualimap [65]. In addition, we checked for any potential large scale mis-assemblies using Sniffles v1.0.8 [66] by mapping the PacBio subreads using NGMLR v0.2.6 [66]. SPAdes v3.7.1 [67] was run on the Illumina data to detect smaller plasmids that might have been lost in the size selection step. The genome was annotated with the NCBI’s prokaryotic genome annotation pipeline (v3.3) [68]. Prophages were identified with Phaster [69]. Detailed annotations for all CDS were computed as described previously [70]; this included assignment to Cluster of Orthologous Groups (COG) categories using eggnog-mapper (v 1.0.3) and EggNOG 4.5, an Interproscan analysis and prediction or /integration of subcellular localizations, lipoproteins, transmembrane helices and signal peptides (for details, see Supplementary **Table 2**).

### Comparative genomics of selected PAO1 genomes

The genome of the *P. aeruginosa* PAO1-UW reference strain [2] was compared to our complete MPAO1 genome using the software Roary (v3.8.0) [71] to define core and strain-specific gene clusters as described before [30, 71]. A BlastP analysis helped to correctly identify conserved genes with ribosomal slippage (*prfB*; *p*eptide chain release factor B) or that encode a selenocysteine (MPAO1_25645), which otherwise can be misclassified as unique genes; genes of 120 bp or below (17 in MPAO1) were not considered. ProgressiveMauve [72] was used to align the genomes globally and to identify larger genomic differences. Smaller differences (indels, single nucleotide polymorphisms (SNPs)), were identified and annotated against the PAO1 reference strain as described previously [70]. Furthermore, contigs from the MPAO1/P1 genome [32] were aligned to our complete MPAO1 genome assembly using BWA mem [73]. Bedtools v2.16.1 ‘genomecov’ [74] was used to calculate a gene-wise coverage, allowing to identify genes that were missed in the 140 contigs.

### Re-mapping of Transposon sequencing data

MPAO1 Tn-seq datasets [24] were downloaded from NCBI’s SRA (SRP052838) and mapped back both to the PAO1-UW reference strain genome [2] and to our MPAO1 assembly following the scripts and notes provided in the Supplement. Insertion sites were computed as described by the authors, reads mapping to multiple genome positions were assigned randomly, and the number of insertion sites per gene was used to differentiate essential and non-essential genes as described [24]. Genes with zero insertions were considered essential; for the remaining genes, normalized read counts across all insertion sites per gene (considering insertions falling within 5-90% of the length of each gene) were log2 transformed and fitted to a normal distribution. Genes with a *p* value < 0.001 were added to the list of essential genes. Finally, essential and conditionally essential genes were identified among the three main growth conditions (LB medium, minimal medium, sputum) as described [24]. Data from each growth condition consisted of multiple mutant pools; for LB, two mutant pools additionally contained multiple replicates (LB-1: 3 replicates; LB-2: 2 replicates). For LB, genes were considered essential in the mutant pool LB-1 and LB-2 if at least two of three (LB-1) and one of two replicates (LB-2) agreed. Next, a consensus set of essential genes in LB and minimal medium was derived from those genes that were essential in at least two of three mutant pools (LB-1, LB-2 and LB-3) in LB and minimal respectively. Similarly, essential genes in sputum (four mutant pools) were derived if data from at least three of four pools agreed. Finally, genes that were essential in all three growth conditions were called “general essential genes (312)” and genes essential to a specific growth condition were called “condition specific essential genes”. Together, they comprise “all essential genes (577)”; for further details, see Supplement.

### Microfluidic chamber used for biofilm growth

The standardized microfluidic flow chamber consisted of a PDMS chip with a straight microfluidic channel (30 mm length × 2 mm width × 0.200 mm depth) that naturally adhered onto a glass coverslip (26 × 60 mm; thickness no.1). The wafer master was fabricated using SU-8 spin-coated to a thickness of 200 μm on a silicon wafer in advance of standard soft lithography replication into PDMS [84]. From this, polyurethane clones of the structures were prepared to upscale production and for sharing microfluidic molds between laboratories. A degassed 10:1 mixture of Sylgard 184 PDMS base and curing agent were cured in an oven at 60°C for 2 h. Following cooling and retrieval from the SU-8 wafer the structured PDMS was attached, structures facing upwards, to a silicone baking mold using transparent double-sided adhesive (3M). The PDMS part was degassed, while the two-component polyurethane (Smooth-Cast™ 310) solutions were each thoroughly shaken for 10 min and then combined in a 1:1 ratio followed by thorough mixing (by repeat inversion and then shaking). The PDMS device was then submerged in the mixture, with degassing for 10 min, after which the mold was left overnight in a well ventilated area followed by a hard bake at 60°C for 4 h. Once cooled the PDMS device was retrieved leaving the polyurethane mold in readiness for replica molding fresh PDMS devices again at 60 °C for 2 h. Importantly, PDMS devices are only retrieved after the polyurethane mold has cooled to room temperature to allow the repeated replication (>100 times) of precision PDMS microfluidic chambers. Inlet and outlet ports were prepared using 1-mm-diameter biopsy punches (Miltex™) and then the device was enclosed using a coverslip that was cleaned with 2% RBS 35 detergent (prepared in demineralized water), rinsed with tap water, then immersed in 96% ethanol and sonicated for 5 min, followed by a final rinse with demineralized water and then autoclaved. The inlet and outlet of the microfluidic flow chamber were connected to a syringe pump with a 25G needle and waste container, respectively, via sterile PTFE tubing (Smiths Medical, ID 0.38 mm, OD 1.09 mm). The chamber was first disinfected by flowing 70% ethanol for 15 min at a rate of 20 μL/min, before rinsing with sterile PBS for 15 min and then flushing with M9 minimal medium (Formedium Ltd, Hunstanto, England) for another 15 min at the same flow rate.

### Device inoculation, biofilm staining and confocal laser microscopy

*P.* aeruginosa MPAO1 was inoculated with 500 µL of an M9-grown overnight pre-culture and grown for ∼16-18 h in 10 mL M9 medium (1x M9 salts supplemented with 2 mM MgSO_4_, 100 μM CaCl_2_ and 5 mM glucose) at 37 °C with gentle rotation (150 rpm) until a cell concentration of 1.5 x 10^9^ bacteria/mL was reached. One mL of the culture was then washed twice with PBS (pH 7.0) by centrifugation at 5,000 xg for 5 min at 10°C. The bacterial pellet was re-suspended and diluted in PBS + 2% M9 such that the final cell suspension contained 3 x 10^8^ bacteria/mL. The microfluidic chamber was set on a hotplate at 37 °C with the glass coverslip in direct contact with the hotplate surface. Freshly prepared bacterial suspension was flown through at a rate of 5 μL/min for 1 h. After 1 h, the bacterial suspension was replaced by M9 medium and run through the system at 5 μL/min for 72 h. After 72 h, CLSM images were taken. The biofilm was stained by flowing 1 mL of Live/Dead (Life Technologies, Oregon, USA) staining solution (1.5 μL Syto9 + 1.5 μL propidium iodide in 1 mL of sterile demineralized water) through the flow chamber at 5 μL/min. Once the channel was filled, the flow was stopped and the biofilm kept in the dark for 30 min to allow dye penetration. Finally, PBS was flown through the system at 5 μL/min for 30 min to remove the staining agent. Confocal imaging was performed using a Leica SP8 with x63 oil immersion lens (HC PL APO CS2 63x/1.30, Southampton; LabA), a Leica SP8 with x63 water immersion lens (HC PL APO 63x/1.20W CORR CS2; BAM, LabB), and a Leica SP2 with x63 water lens (HCX APO L 63x/0.9W; Groningen, LabC) for 3 biological repeats comprising 3 technical repeats per site (n=9 biological / n=27 technical). Z-stacks (1 μm) were taken of the biofilms formed on the PDMS surface of the device at five separate regions (beside the inlet, 25%, 50%, 75%, and beside the outlet). COMSTAT 2.1 (Image J) analysis of combined confocal data was performed to provide a quantification of average biofilm thickness and Live/Dead biovolume [75]. A 2-way ANOVA multiple comparison was performed with Tukey’s post hoc test to determine 95% confidence intervals. Similar conditions were applied to strain PA4726 (*cbrB*) that had shown reduced biofilm growth during screening, and PA3552 (*arnB*) which demonstrated robust biofilm formation. Biofilm formation of both mutant strains was compared to the MPAO1 WT strain after 18 h growth in the flow chamber.

### Screening the public MPAO1 transposon library for antibiotic resistance

The protocol to assess the antibiotic resistance of biofilm-forming MPAO1 cells was adapted from a previous study [76]. Frozen mutant stocks of 50 randomly selected mutants of the UW Genome Center’s *P. aeruginosa* PAO1 transposon mutant library [21], each harboring a transposon insertion inactivating the function of the respective gene, were allowed to recover in 20% BHI overnight at 150 rpm and 37°C. All subsequent incubations were done at 37°C in 96 well plates covered with an air-permeable foil without further shaking. The overnight cultures were diluted 10 fold in M9 medium and 100 µL each was distributed in six plates (1 well/mutant/plate). After 24 h incubation, the biofilm formation from two plates was quantified by crystal violet staining, while biofilms from the other four plates were washed with 0.9% NaCl to remove planktonic bacteria. Bacteria were then exposed to either M9 or M9 supplemented with 25 µg/mL of colistin, i.e., much higher than the minimal inhibitory concentration (MIC) for planktonic growth of *P. aeruginosa* (4 µg/mL), allowing us to focus specifically on the biofilm bacteria. After 24h treatment, the medium was removed, biofilms were washed with 0.9% NaCl to remove all traces of antibiotics, and bacteria were allowed to recover in fresh colistin-free M9 medium. After 24 h incubation, the recovery of biofilm bacteria was measured by turbidity (OD600) to reveal if the mutation influences the resistance attributed by the biofilm. To confirm the reliability of our screening, promising mutants were analyzed independently in triplicate. Cell suspensions of each mutant were prepared in M9 medium (5 x10^6^ CFU/mL) and biofilm biomass was quantified by crystal violet after 24h incubation at 37°C. Biofilm cells resistance was quantified by measuring the turbidity of biofilm suspension after 24h treatment with different concentrations of colistin and after 24h recovery in fresh M9 medium.

### Protein extraction from MPAO1 planktonic and biofilm cultures

For planktonic protein extractions, 10 mL MPAO1 was grown overnight (∼18 h) in M9 medium under gentle rotation (150 rpm), centrifuged at 4,000 xg/5 min/RT, and the pellet resuspended in 1 mL Hanks’ Balanced Salt Solution (HBSS). Biofilms were grown for 72 h using the microfluidic device as previously described, the PDMS device removed from the glass coverslip, and the combined biofilm biomass from 3 lanes harvested into 1 mL HBSS. Cells from both populations were washed twice in HBSS at 10,000 xg/5 min/RT and the pellets resuspended in 1 mL lysis buffer (7 M urea, 2 M thiourea, 35 mM CHAPS, 20 mM DTT, 1 M NaCl). Samples were frozen at -80 °C for 30 min and then thawed at 34 °C for 20 min. Trichloroacetic acid (TCA) precipitation was performed by adding the bacterial samples to 100% ice-cold acetone and 100% trichloroacetic acid in a 1:8:1 ratio and precipitating at -20 °C for 1 h. Samples were then centrifuged (18,000 xg/10 min/4 °C), the supernatant discarded, and the pellet washed twice with 1 mL ice-cold acetone (18,000 xg/10 min/4 °C). Acetone was removed, the pellet air-dried at room temperature, and resuspended in 0.1 M Triethylammonium bicarbonate (TEAB) plus 0.1 % Rapigest. Protein sample validation was performed by 1DE gel electrophoresis. 19.5 μL sample was added to 7.5 μL NuPAGE LDS buffer and 3 μL NuPAGE reducing reagent, heated at 70 °C for 10 min, then run on a NuPAGE 4-12% Bis-Tris gel with MOPS buffer at 200 V for 50 min alongside a Novex Sharp standard. The gel was stained with SimplyBlue Safe Stain for 1 h, then destained with dH_2_O.

### Protein processing, mass spectrometry and database search

Protein samples were heated at 80 °C for 10 min, then DTT added at a final concentration of 2 mM and incubated at 60 °C for 45 min. Samples were then briefly vortexed, pulse spun, and cooled to room temperature before adding iodoacetamide to a final concentration of 6 mM. Samples were incubated at room temperature for 45 min (protected from light), vortexed and pulse spun briefly, then trypsin added at a final concentration of 1.3 µg/mL. Following incubation overnight at 37 °C (protected from light), trifluoroacetic acid (TFA) was added to a final concentration of 0.5% then incubated at 37 °C for 30 min. Samples were centrifuged at 13,000 xg for 10 min at RT, the supernatants removed and vacuum concentrated. The resultant pellets were resuspended in 3% acetonitrile + 0.1% trifluoroacetic acid and peptide quantification performed using the Direct Detect system (Merck Millipore). Protein samples were normalized then vacuum concentrated in preparation for mass spectrometry.

Peptide extracts (1 μg on column) were separated on an Ultimate 3000 RSLC nano system (Thermo Scientific) using a PepMap C18 EASY-Spray LC column, 2 μm particle size, 75 μm x 75 cm column (Thermo Scientific) over a 140 min (single run) linear gradient of 3–25% buffer B (0.1% formic acid in acetonitrile (v/v)) in buffer A (0.1% formic acid in water (v/v)) at a flow rate of 300 nL/min. Peptides were introduced using an EASY-Spray source at 2000 V to a Fusion Tribrid Orbitrap mass spectrometer (Thermo Scientific). The ion transfer tube temperature was set to 275 °C. Full MS spectra were recorded from 300 to 1500 *m/z* in the Orbitrap at 120,000 resolution using TopSpeed mode at a cycle time of 3 s. Peptide ions were isolated using an isolation width of 1.6 amu and trapped at a maximal injection time of 120 ms with an AGC target of 300,000. Higher-energy collisional dissociation (HCD) fragmentation was induced at an energy setting of 28 for peptides with a charge state of 2–4. Fragments were analysed in the Orbitrap at 30,000 resolution. Analysis of raw data was performed using Proteome Discoverer software (Thermo Scientific) and the data processed to generate reduced charge state and deisotoped precursor and associated product ion peak lists. These peak lists were searched against the *P. aeruginosa* MPAO1 protein database (a max. of one missed cleavage was allowed for tryptic digestion, variable modification was set to contain oxidation of methionine and N-terminal protein acetylation, and carboxyamidomethylation of cysteine was set as a fixed modification). The FDR was estimated with randomized decoy database searches and was filtered to below 1% FDR at the protein level. Differentially expressed proteins were identified using DESeq2 [77]; significantly differentially expressed proteins had an adjusted *p* value ≤ 0.05 and a log_2_ fold change of ≥ 1 or ≤ -1.

### Proteogenomics

An iPtgxDB was created for *P. aeruginosa* MPAO1 as described previously [31], using the NCBI annotation as anchor annotation. *Ab initio* gene predictions from Prodigal [78] and ChemGenome [79] and a modified *in silico* prediction that considers alternative start codons (TTG, GTG, CTG) and ORFs above 6 amino acids (aa) in length were integrated in a step-wise fashion. Proteomics data from MPAO1 cells grown planktonically or as biofilm were searched against this iPtgxDB with MS-GF+ (v2019.04.18) [80] using Cysteine carbamidomethylation as fixed, and oxidation of methionine as variable modifications. Using the target-decoy approach of MS-GF+, the FDR at the PSM level was estimated and filtered below 0.2%. Only unambiguous peptides as identified by a PeptideClassifier analysis [48], using the extension that supports proteogenomics for prokaryotes [31], were considered.

## Acknowledgements

The authors thank Jürg Frey and Daniel Frei (Agroscope) for generating Illumina MiSeq data. The authors acknowledge funding from the Joint Programming Initiative against AntiMicrobial Resistance (JPIAMR) and national grants to JSW for RNA (MRC MR/R005621/1), HvdM for OCO (ZonMW grant #547001003), QR for JV and MTB (SNSF grant 40AR40_173611), and to FS for FP (BMBF #01KI1710). CHA acknowledges funding for ARV through grants 156320 and 188722 from the SNSF.

## Author contributions

VS and ARV carried out genome assembly. ARV performed comparative genomics analyses, remapped existing Tn-seq data, created the iPtgxDB, performed proteogenomics analyses and created figures with CHA. MTB grew cells and extracted gDNA. JV devised and carried out the screening approach, overseen by QR. OCO and HvdM designed the mold for the microfluidic flow chamber and JW provided device replication expertise. Microfluidic-confined biofilm growth and confirmed reproducibility were undertaken by RNA, JSW, OCO, HvdM, FP and FS. PS generated shotgun proteomics data from planktonic and biofilm cells provided by RNA. RNA and JSW carried out biofilm growth in the mold for selected mutants from the transposon mutant collection and analyzed proteomics data. CHA oversaw genome sequencing and assembly, comparative genomics and proteogenomics, and wrote the manuscript together with input from RNA, FS, ARV and all other authors.

## Data Access

The MPAO1 genome sequence is available at NCBI Genbank (acc# CP027857; Bioproject: PRJNA438597, Biosample: SAMN08722738). Read data are available under SRR10153205 (Illumina) and SRR10153206 (PacBio). Proteomics data are available from PRIDE (acc# PXD017122) upon acceptance of the manuscript. The iPtgxDB for *P. aeruginosa* MPAO1 is available from https://iptgxdb.expasy.org, both as a searchable protein database (FASTA format) and a GFF file, which can be loaded in a genome viewer and overlaid with experimental evidence. Biofilm growth data from the microfluidic chamber is available at https://metafluidics.org/device-keywords/microbiology/. To support technology dissemination, the polyurethane master molds of the microfluidic chambers are available upon request from the UoS/NBIC; a CAD file can be found as Supplementary **File 11**.

