## Supplemental Information for "An integrated model system to gain mechanistic insights into biofilm formation and antimicrobial resistance development in *Pseudomonas aeruginosa* MPAO1"

### Supplemental Material

#### Table of Contents

**Table S1. Genome characteristics of *P. aeruginosa* MPAO1 and PAO1-UW.** 2
**Table S2. Detailed annotation & integrated information for data mining (Master table).** 3
**Figure S1. The genome of strain MPAO1/P1 contains more interrupted genes.** 4
**Table S3. Summary of SNP differences between strain MPAO1 and PAO1.** 5
**Table S4. List of shared and specific gene clusters for strains PAO1 and MPAO1.** 5
**Table S5. Gene ontology categories among 232 unique MPAO1 gene clusters.** 6
**Table S6. Re-mapping of Tn-seq data against the MPAO1 genome leads to a higher percentage of mapped reads.** 7
**Figure S2. Overview of conditionally essential genes in *P. aeruginosa* MPAO1.** 8
**Table S7. Summary table of 1117 genes essential in at least one Tn-seq library.** 8
**Table S8. Laminar flow conditions achieved in the biofilm chamber.** 9
**Table S9. List of *P. aeruginosa* MPAO1 mutant strains used in this work.** 10
**Figure S3. Several members of the H1-T6SS are upregulated in biofilm.** 12
**Table S10. Summary of annotation clusters created for the MPAO1 iPtgxDB.** 13
**File S11. Design of the microfluidic chamber as a CAD file.** 13
**References** 14

**Table S1. Genome characteristics of *P. aeruginosa* MPAO1 and PAO1-UW.**

| | ***P. aeruginosa* MPAO1** | ***P. aeruginosa* PAO1-UW*** |
|---|---|---|
| Genbank accession # | CP027857 | NC_002516 |
| No. chromosomes (plasmids) | 1 (0) | 1 (0) |
| Size (bp) | 6,275,467 | 6,264,404 |
| G+C content (%) | 66.5 | 66.6 |
| Coverage (PacBio) | 180x | n.a. |
| Coverage (Illumina MiSeq) | 101x | n.a. |
| Total No. of genes | 5,926 | 5,697 |
| No. of protein-coding genes (CDSs) | 5,799 | 5,572 |
| No. of rRNA operons (16S, 23S, 5S) | 4,4,4 | 4,4,4 |
| No. of tRNA genes | 63 | 63 |
| No. of pseudogenes | 48 | 19 |
| No. of ncRNA, tmRNA | 4, - | 29, 1 |
| No. of 4.5S rRNA | - | 1 |
| Prophages | 3 | 2 |

*We interchangeably use *P. aeruginosa* PAO1 or PAO1-UW.

**Table S2. Detailed annotation & integrated information for data mining (Master table).**

See separate Excel file. The file contains detailed annotation for all 5,799 annotated MPAO1 CDS. Furthermore, it includes information whether the genes are conserved or unique compared to *P. aeruginosa* PAO1, the respective PAO1 homolog (and gene name, where applicable), the genes missed in three Illumina short read-based assemblies of MPAO1 strains [1] [2], gene essentiality status from our study and previous data sets [3] [4], protein expression evidence from our study (biofilm versus planktonic growth), and additional information about protein domains, families, patterns, signatures, a Gene Ontology (GO) classification, a prediction of subcellular localization, lipoproteins, etc. We did not specifically assess short MPAO1-unique genes, whose gene essentiality status is more difficult to robustly classify. Instead, we added a proteogenomics element to enable identification of novel short proteins (main article).

**Figure S1**. **The genome of strain MPAO1/P1 contains more interrupted genes.**

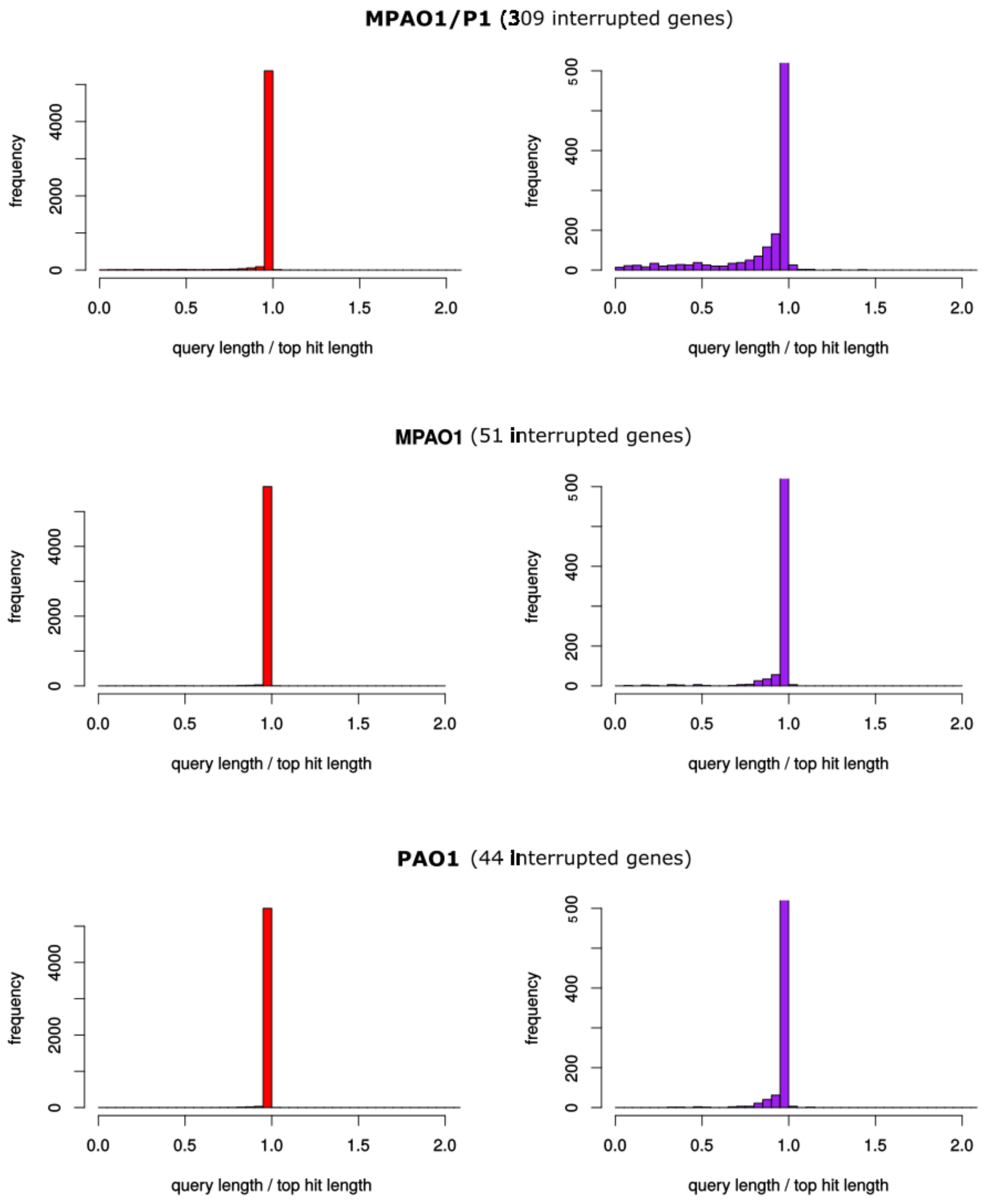


An analysis with Ideel (https://github.com/mw55309/ideel) uncovered large differences of the number of predicted pseudogenes/interrupted genes, which can serve as one parameter to estimate genome completeness. For MPAO1/P1 (5,791 CDS), about six times as many putative pseudogenes/interrupted genes were identified compared to the complete MPAO1 (5,799 CDS) and PAO1-UW (5,572 CDS) genome sequences (MPAO1/P1: 309; MPAO1: 51; PAO1-UW: 44). A complete genome typically shows a narrow peak around 1, i.e., most of the CDS have a full length BlastP hit against the respective UniProt entry, and a shallow tail of the distribution towards the left (see zoomed region in the right plots).

**Table S3. Summary of SNP differences between strain MPAO1 and PAO1.**

The table (see separate Excel file) lists the differences between our complete genome assembly of MPAO1 and the genome sequence of PAO1-UW [5], the PAO1 type strain. We could confirm the 16 SNPs reported previously for both strains [6], one SNP only in MPAO1 and six SNPs observed as base exchanges in both strains. We could also confirm nine of the SNPs that had been reported for the PAO1 DSM strain only [6] (synonymous substitutions in Phage pf1 protein) which are located at the beginning of the inversion region, one SNP in transcriptional regulator MexT and one intergenic SNP at position 5,033,102 of the MPAO1 genome. In addition, we observed a total of 176 additional SNPs and INDELs between PAO1 and MPAO1 that were not reported by Klockgether and colleagues, as their comparison had focused on selected genomic regions of the two strains [6].

**Table S4. List of shared and specific gene clusters for strains PAO1 and MPAO1.**
Gene clusters specific to either PAO1 (21) or MPAO1 (232), and those shared between the two strains (5,534) as returned from an analysis with Roary [7] are listed in a separate Excel file, along with genomic coordinates, annotation and COG classification. For the MPAO1-specific gene clusters, information about essentiality was computed based on the very important dataset by Lee and colleagues and using the scripts they provided in their Supplementary Material [3].

**Table S5. Gene ontology categories among 232 unique MPAO1 gene clusters.**

| GO accession | GO description | Unique genes | total proteins annot. | unique proteins annot. | p-value |
|---|---|---|---|---|---|
| **Biological Process** | | | | | |
| GO:0006468 | protein phosphorylation | MPAO1_11765, MPAO1_24875, MPAO1_24880 | 10 | 3 | 2.90E-05 |
| GO:0030153 | bacteriocin immunity | MPAO1_05695, MPAO1_20105 | 5 | 2 | 0.00041 |
| GO:0006571 | tyrosine biosynthetic process | MPAO1_09400 | 1 | 1 | 0.00664 |
| GO:0006470 | protein dephosphorylation | MPAO1_24870 | 5 | 1 | 0.03276 |
| GO:0015074 | DNA integration | MPAO1_24800 | 7 | 1 | 0.04557 |
| **Molecular Function** | | | | | |
| GO:0004672 | protein kinase activity | MPAO1_11765, MPAO1_24875, MPAO1_24880 | 85 | 3 | 0.0226 |
| GO:0003866 | 3-phosphoshikimate 1-carboxyvinyltransferase | MPAO1_09400 | 1 | 1 | 0.0073 |
| GO:0004308 | exo-alpha-sialidase activity | MPAO1_11350 | 1 | 1 | 0.0073 |
| GO:0004665 | prephenate dehydrogenase (NADP+) activity | MPAO1_09400 | 1 | 1 | 0.0073 |
| GO:0008849 | enterochelin esterase activity | MPAO1_13245 | 1 | 1 | 0.0073 |
| GO:0008977 | prephenate dehydrogenase (NAD+) activity | MPAO1_09400 | 1 | 1 | 0.0073 |
| GO:0008998 | ribonucleoside-triphosphate reductase activity | MPAO1_16050 | 2 | 1 | 0.0145 |
| GO:0004722 | protein serine/threonine phosphatase activity | MPAO1_24870 | 3 | 1 | 0.0217 |
| GO:0015643 | toxic substance binding | MPAO1_20105 | 3 | 1 | 0.0217 |
| GO:0005102 | receptor binding | MPAO1_24810 | 4 | 1 | 0.0288 |
| GO:0016805 | dipeptidase activity | MPAO1_06140 | 4 | 1 | 0.0288 |

GO categories for which only one gene among the unique MPAO1 genes was affected are shown in gray

**Table S6. Re-mapping of Tn-seq data against the MPAO1 genome leads to a higher percentage of mapped reads.**

| Library Name | # mapped reads | | % increase mapped reads | # unique insertion sites | | % increase unique insertion sites |
|---|---|---|---|---|---|---|
| | PAO1-UW | MPAO1 | | PAO1-UW | MPAO1 | |
| LB-1_Rep1 | 2,095,460 | 2,099,090 | 0.17% | 92,731 | 92,938 | 0.22% |
| LB-1_Rep2 | 13,409,799 | 13,427,972 | 0.14% | 89,678 | 89,884 | 0.23% |
| LB-1_Rep3 | 8,970,876 | 8,990,866 | 0.22% | 64,155 | 64,298 | 0.22% |
| LB-2_Rep1 | 14,495,458 | 14,546,760 | 0.35% | 82,060 | 82,239 | 0.22% |
| LB-2_Rep2 | 18,630,794 | 18,648,695 | 0.10% | 82,441 | 82,655 | 0.26% |
| LB_3 | 23,922,898 | 23,952,863 | 0.13% | 110,102 | 110,331 | 0.21% |
| Minimal-1 | 27,740,278 | 27,770,384 | 0.11% | 193,648 | 194,051 | 0.21% |
| Minimal-2 | 13,365,514 | 13,383,055 | 0.13% | 204,750 | 205,193 | 0.22% |
| Minimal-3 | 6,441,754 | 6,449,398 | 0.12% | 250,831 | 251,570 | 0.29% |
| Sputum-1 | 5,630,125 | 5,635,321 | 0.09% | 78,706 | 78,921 | 0.27% |
| Sputum-2 | 7,609,070 | 7,617,184 | 0.11% | 181,365 | 181,737 | 0.21% |
| Sputum-3 | 14,491,394 | 14,508,287 | 0.12% | 48,809 | 48,930 | 0.25% |
| Sputum-4 | 2,991,136 | 2,994,255 | 0.10% | 91,856 | 92,035 | 0.19% |
| HumanSerum | 754,467 | 755,143 | 0.09% | 127,543 | 127,808 | 0.21% |
| 0.1XLB | 8,735,565 | 8,743,902 | 0.10% | 121,430 | 121,679 | 0.21% |
| BHI | 4,438,845 | 4,443,736 | 0.11% | 174,419 | 174,796 | 0.22% |

As expected, consistently higher percentages of mapped reads were achieved when mapping the Tn-seq datasets to the complete genome assembly of *P. aeruginosa* strain MPAO1 compared to mapping it to the reference strain PAO1-UW.

**Figure S2. Overview of conditionally essential genes in *P. aeruginosa* MPAO1.**

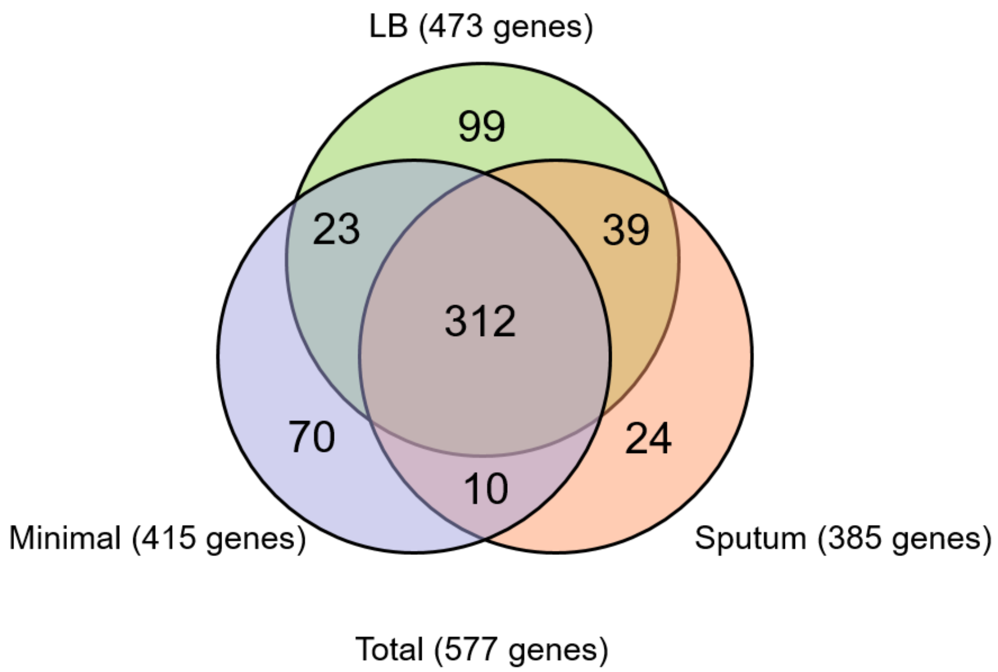


Using the scripts released by Lee and colleagues [3], Tn-seq data were re-mapped to our complete MPAO1 genome and compared with data published, i.e., the set of genes essential in either one of the three conditions sputum, minimal medium and LB medium (577 "all essential genes"), as well as the 312 genes essential in all 3 categories ("general essential genes"). This analysis was close to the original results. Due to the higher mapping success of reads to the complete genome sequence, we identified 39 MPAO1-unique "all essential genes" in the MPAO1 genome (Table 2 and Table S5). Overall, 1117 genes were identified as essential in at least one Tn-seq library; thereof, 136 were MPAO1-unique genes (see Table S7). Six of the MPAO1-unique genes were general essential genes.

**Table S7. Summary table of 1117 genes essential in at least one Tn-seq library.**

See separate Excel file. For completeness, we also show the MPAO1-unique genes that were identified as essential in at least one of the sixteen Tn-seq samples (136; Table S7). The second column shows in how many of the 16 samples a respective gene was called essential. The last three columns indicate the subset of 577 genes essential in at least one of the three primary growth conditions (LB, minimal and sputum), and the subset of 312 genes essential in all three primary growth conditions, i.e., general essential genes (see Methods).

**Table S8. Laminar flow conditions achieved in the biofilm chamber.**

The flow in the flow chamber had a defined laminar flow. The calculated Reynolds Number (see formulas below table) was 0.103 given a volumetric flow rate 5 µL/min, pressure 0.0108 mbar, the channel specifications given below, and correcting for the viscosity and density of water at 37 °C.

| **Initial Conditions** | |
|---|---|
| Channel geometry | rectangular |
| Channel height | 200 µm |
| Channel width | 2000 µm |
| Channel length | 30 mm |
| Fluid viscosity | 0.73 cP (Pa.s / $10^3$) |
| Fluid density | 0.993 g/cm$^3$ (kg/m$^3$ x$10^3$) |
| **Results** | |
| Pressure | 0.0107652 mbar |
| Flow rate | 5 µL/min |
| Velocity | 0.000208333 |
| Reynolds number | 0.103051 (**laminar flow**) |

$$Re = \frac{d * D * v}{\mu}$$

Re: Reynolds Number
d: Density
v: Flow Velocity
D: Hydraulic diameter
µ: Dynamic Viscosity

$$D = \frac{4A}{P}$$

A: section area

P: wettered perimeter

**Table S9. List of *P. aeruginosa* MPAO1 mutant strains used in this work.**

All mutants are from the UW Genome Center *P. aeruginosa* MPAO1 transposon mutant library (laboratory of Prof. Dr. Colin Manoil). Listed are the name of the mutant strain, the identifier of the respective gene in PAO1-UW and the MPAO1 locus tag, the gene name and a putative function. Note: gene names in the UW library are listed as PA0160-G03::ISphoA/hah (given here for strain PW1274 as one example).

| Strain name | PAO1 gene identifier | MPAO1 locus tag | Gene name | Putative function |
|---|---|---|---|---|
| PW1274 | **PA0160** | MPAO1_00860 | | hypothetical protein |
| PW7893 | **PA0357** | MPAO1_01890 | *mutM* | formamidopyrimidine-DNA glycosylase |
| PW1808 | **PA0440** | MPAO1_02325 | | probable oxidoreductase |
| PW1871 | **PA0476** | MPAO1_02520 | | probable permease |
| PW2290 | **PA0711** | MPAO1_22480 | | hypothetical protein |
| PW2385 | **PA0761** | MPAO1_22195 | *nadB* | L-aspartate oxidase |
| PW2642 | **PA0898** | MPAO1_21485 | *aruD* | succinylglutamate 5-semialdehyde dehydrogenase |
| PW2661 | **PA0914** | MPAO1_21390 | | hypothetical protein |
| PW3211 | **PA1224** | MPAO1_19730 | | probable NAD(P)H dehydrogenase |
| PW3497 | **PA1373** | MPAO1_18945 | *fabF2* | 3-oxoacyl-acyl carrier protein synthase II |
| PW3660 | **PA1467** | MPAO1_18470 | | hypothetical protein |
| PW3859 | **PA1599** | MPAO1_17765 | | probable transcriptional regulator |
| PW3904 | **PA1629** | MPAO1_17615 | | probable enoyl-CoA hydratase/isomerase |
| PW4005 | **PA1693** | MPAO1_17275 | *pscR* | translocation protein in type III secretion |
| PW4095 | **PA1755** | MPAO1_16955 | | hypothetical protein |
| PW4171 | **PA1804** | MPAO1_16665 | *hupB* | DNA-binding protein HU |
| PW4474 | **PA1997** | MPAO1_15660 | | probable AMP-binding enzyme |
| PW4590 | **PA2084** | MPAO1_15180 | | probable asparagine synthetase |
| PW4975 | **PA2361** | MPAO1_13710 | | hypothetical protein |
| PW5552 | **PA2716** | MPAO1_11820 | | probable FMN oxidoreductase |
| PW5732 | **PA2825** | MPAO1_11185 | | probable transcriptional regulator |
| PW5923 | **PA2928** | MPAO1_10660 | | hypothetical protein |
| PW6141 | **PA3064** | MPAO1_09940 | *pelA* | PelA |
| PW6275 | **PA3137** | MPAO1_09535 | | probable major facilitator superfamily (MFS) transporter |
| PW6504 | **PA3279** | MPAO1_08785 | *oprP* | Phosphate-specific outer membrane porin OprP precursor |
| PW6719 | **PA3391** | MPAO1_08185 | *nosR* | regulatory protein NosR |
| PW6755 | **PA3410** | MPAO1_08085 | | probable sigma-70 factor, ECF subfamily |
| PW6868 | **PA3470** | MPAO1_07770 | | hypothetical protein |
| PW6985 | **PA3534** | MPAO1_07440 | | probable oxidoreductase |

| | | | | |
|---|---|---|---|---|
| PW7021 | **PA3552** | MPAO1_07345 | *arnB* | ArnB |
| PW7067 | **PA3574** | MPAO1_07230 | | probable transcriptional regulator |
| PW7383 | **PA3772** | MPAO1_06190 | | hypothetical protein |
| PW7566 | **PA3890** | MPAO1_05560 | | probable permease of ABC transporter |
| PW8169 | **PA4224** | MPAO1_03835 | *pchG* | pyochelin biosynthetic protein PchG |
| PW8212 | **PA4282** | MPAO1_22735 | | probable exonuclease |
| PW8322 | **PA4338** | MPAO1_23025 | | hypothetical protein |
| PW8365 | **PA4362** | MPAO1_23150 | | hypothetical protein |
| PW8707 | **PA4578** | MPAO1_24275 | | hypothetical protein |
| PW8936 | **PA4711** | MPAO1_25105 | | hypothetical protein |
| PW8965 | **PA4726** | MPAO1_25185 | *cbrB* | two-component response regulator CbrB |
| PW9431 | **PA5020** | MPAO1_26730 | | probable acyl-CoA dehydrogenase |
| PW9793 | **PA5219** | MPAO1_27780 | | hypothetical protein |
| PW9856 | **PA5261** | MPAO1_28000 | *algR* | alginate biosynthesis regulatory protein AlgR |
| PW9891 | **PA5281** | MPAO1_28105 | | probable hydrolase |
| PW9895 | **PA5283** | MPAO1_28115 | | probable transcriptional regulator |
| PW9934 | **PA5304** | MPAO1_28225 | *dadA* | D-amino acid dehydrogenase, small subunit |
| PW10082 | **PA5384** | MPAO1_28660 | | probable lipolytic enzyme |
| PW10195 | **PA5442** | MPAO1_28970 | | conserved hypothetical protein |
| PW10219 | **PA5455** | MPAO1_29035 | | hypothetical protein |

**Figure S3**. **Several members of the H1-T6SS are upregulated in biofilm.**

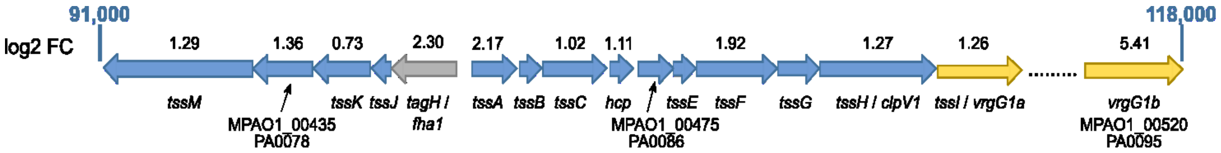


Genomic region of *P. aeruginosa* MPAO1 that shows part of the H1-T6SS region (roughly from nucleotide 91,000-118,000); gene names or MPAO1 locus tags are shown (with the respective PAO1 homolog below). The colors were selected as in [8] and indicate structural elements (blue), vgrGs (yellow) and other known T6SS genes (gray). The shotgun proteomics data (log2 fold change biofilm over planktonic is shown above the arrows) indicated that several (9 of 14, 64%) of the proteins encoded by structural elements of the H1-T6SS and secreted proteins [9] were upregulated in biofilm cells compared to planktonic cells. The H1-T6SS has been described as "a molecular gun firing toxins (Tse1-Tse7)" and has been implied "to challenge the survival of other bacteria and help *P. aeruginosa* prevail in specific niches" [8]. This specific H1-T6SS has recently been shown to be highly relevant for the ability of *P. aeruginosa* strains to dominate in multi-species biofilms [10]. Notably, all three VgrG proteins that are co-regulated with this T6SS (VgrG1a-c) were upregulated, including VGR1c (MPAO1_11985; PA2685; 2.53 log2 FC). In contrast, none of the seven other members of the VgrG family (total of 10) [9] was expressed (see Supplementary **Table 2**).

Our pilot study proteomics dataset covered about 33% of the annotated MPAO1 proteins. This coverage was below that of the extensive proteomics dataset that had allowed to uncover expression evidence for all *Bartonella henseale* Type IV secretion system (T4SS) members [11]. However, that coverage was only be achieved by employing several elaborate fractionation and enrichment strategies, which was beyond the scope of this pilot study. Several of the structural members of the H1-T6SS include shorter proteins and membrane proteins, both of which are more difficult to detect by shotgun proteomics.

**Table S10. Summary of annotation clusters created for the MPAO1 iPtgxDB.**

| Annotation source (tag in identifier) | Anno-tations | Clusters* | New clusters | New reductions | New extensions | Total clusters | Total ids |
|---|---|---|---|---|---|---|---|
| RefSeq (refseq) | 5,851 | 5,851 | 5,851 | 0 | 0 | 5,851 | 5,851 |
| Prodigal (prod) | 5,691 | 5,691 | 55 | 309 | 150 | 5,906 | 6,365 |
| ChemGenome (chemg) | 4,616 | 4,616 | 1,703 | 81 | 1,745 | 7,609 | 9,894 |
| *In silico* ORF (orf) | 155,710 | 70,655 | 63,065 | 78 | 76,881 | 70,674 | 149,918 |

* See the original paper for a detailed description of the annotation clusters [12] or also the website https://iptgxdb.expasy.org/creating_iptgxdbs/ for more information.

**File S11. Design of the microfluidic chamber as a CAD file.**

See separate DWG file.
